# Ferric Heme as a CO/NO Sensor in the Nuclear Receptor Reverbβ by Coupling Gas binding to Electron Transfer

**DOI:** 10.1101/2020.06.22.164806

**Authors:** Anindita Sarkar, Eric L. Carter, Jill B. Harland, Amy L. Speelman, Nicolai Lehnert, Stephen W. Ragsdale

## Abstract

Rev-Erbβ is a nuclear receptor that couples circadian rhythm, metabolism, and inflammation.^1-7^ Heme binding to the protein modulates its function as a repressor, its stability, its ability to bind other proteins, and its activity in gas sensing.^8-11^ Rev-Erbβ binds Fe^3+^-heme tighter than Fe^2+^-heme, suggesting its activities may be regulated by the heme redox state.^9^ Yet, this critical role of heme redox chemistry in defining the protein’s resting state and function is unknown. We demonstrate by electrochemical and whole-cell electron paramagnetic resonance experiments that Rev-Erbβ exists in the Fe^3+^ form within the cell essentially allowing the protein to be heme-replete even at low concentrations of labile heme in the nucleus. However, being in the Fe^3+^ redox state contradicts Rev-Erb’s known function as a gas sensor, which dogma asserts must be a Fe^2+^ protein This paper explains why the resting Fe^3+^-state is congruent both with heme-binding and cellular gas sensing. We show that the binding of CO/NO elicits a striking increase in the redox potential of the Fe^3+^/Fe^2+^ couple, characteristic of an EC mechanism in which the unfavorable ***E***lectrochemical reduction of heme is coupled to the highly favorable ***C***hemical reaction of gas binding, making the reduction spontaneous. Thus, Fe^3+^-Rev-Erbβ remains heme-loaded, crucial for its repressor activity, and only undergoes reduction when diatomic gases are present. This work has broad implications for hemoproteins where ligand-triggered redox changes cause conformational changes influencing protein’s function or inter-protein interactions, like NCoR1 for Rev-Erbβ. This study opens up the possibility of CO/NO-mediated regulation of the circadian rhythm through redox changes in Rev-Erbβ.

## Introduction

Rev-Erb-α and -β belong to the nuclear receptor superfamily. They are indispensable components of the circadian rhythm and regulate the expression of genes involved in lipid and glucose metabolism and inflammatory responses.^1-7,11,12^ Rev-Erbs are also implicated in influencing cognitive and neuronal functions.^13-15^ Nuclear receptors have a characteristic N-terminal AB region followed by a DNA binding C-domain linked to a ligand binding domain (LBD) via a flexible linker.^16^ Heme binding to the LBD promotes proteasomal protein degradation, facilitates its interaction with co-repressors, and enhances its transcriptional repression activity. ^8,9,11,17^

In Rev-Erbβ-LBD, a Cys-Pro heme regulatory motif affords the Cys as an axial ligand to Fe^3+^-heme while a second axial ligation is provided by a distal Histidine (His) residue (Fig. 1a).^18,19^ Besides adopting a His/Cys ligated 6-coordinate low-spin (LS) Fe^3+^ state, Rev-Erbβ can undergo a thiol-disulfide redox switch wherein the coordinating Cys384 forms a disulfide bond with Cys374 giving rise to a His/neutral residue ligated Fe^3+^ heme state.^18,20^ A metal-based redox switch equilibrates Rev-Erbβ between the Fe^3+^ and Fe^2+^ states.^18-20^

**Fig. 1.**
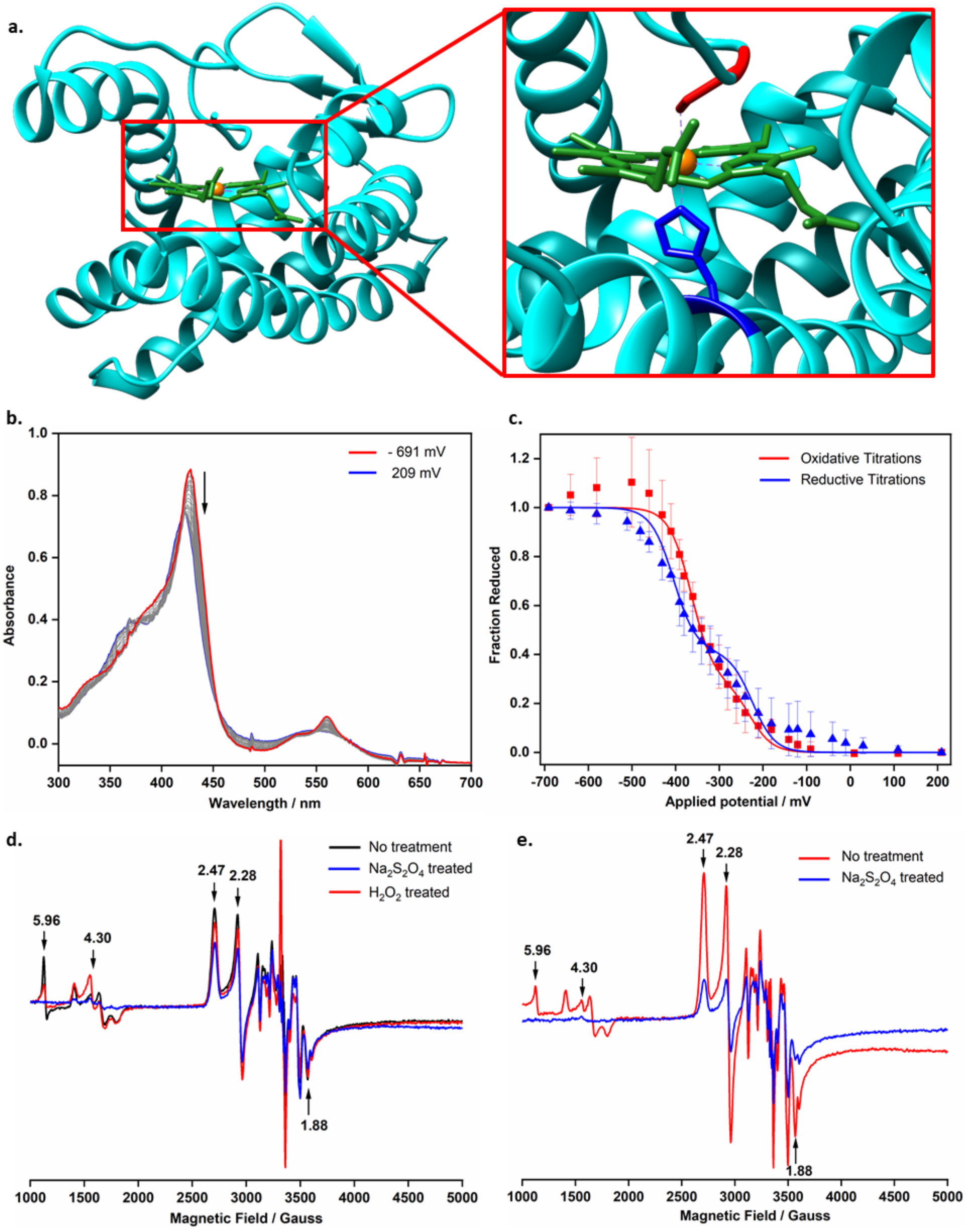
Rev-Erbβ LBD exists in Fe^3+^ not Fe^2+^ form in cellulo. **a**, Crystal structure of Rev-Erbβ LBD complexed with Fe^3+^ heme (3CQV), ligand binding pocket depicting the Fe^3+^ heme coordinated by Cys (red) and His (blue). **b**, Spectral changes observed during potentiometric titration of Rev-Erbβ LBD (82 μM); **c**, Fractional Fe^2+^-protein (calculated from relative absorption at 559 nm) versus applied potential and theoretical Nernst curves for a one-electron redox reaction. Individual data points presented as mean ± s.d. The redox titrations of the LBD are based on intensity changes in the α bands and include 3 and 5 datasets with Soret maxima at 422 nm and 427 nm, respectively; **d**, EPR spectra of *E. coli* cells overexpressing Rev-Erbβ LBD without any treatment and after incubation with with dithionite (20 mM) and H_2_O_2_ (20 mM) for 3 min; **e**, EPR spectra of *E. coli* cells overexpressing Rev-Erbβ LBD without any treatment, and with a pinch of solid dithionite for 5 min. The *g-*values for the rhombic feature of the LBD iron (2.47, 2.28, 1.88), the adventitious iron (4.3) and the HS iron (5.96) are depicted in the EPR spectra. EPR Conditions: temperature 11 K, microwave power 208 µW; microwave frequencies for **(d)** 9.3851 (black), 9.3881 (blue), 9.3902 (red) GHz and for **(e)** 9.3840 (red), 9.3873 (blue) GHz, modulation frequency 100 kHz, modulation amplitude 7 G, 2 scans, 327.68 ms time constant.

The poise between both thiol-disulfide and the Fe^3+/2+^ heme groups of Rev-Erbβ are important in determining the heme occupancy of the protein *in cellulo*, given that the labile heme pools in the nucleus are in low nanomolar concentrations (∼<2.5 nM).^21^ Here we focus on the Fe^3+/2+^ redox couple and how this contributes to Rev-Erb’s ability to be a gas sensor. The heme-binding constants for the different redox states of Rev-Erbβ range over 200-fold, from the Fe^3+^/thiolate with K_d_ ∼ 0.1 nM to Fe^2+^ with K_d_ ∼ 22 nM.^9^ These K_d_ values indicate that Rev-Erb would lose a significant amount of heme if it is reduced to the ferrous form, and thus a role as a gas sensor seems incongruent since the Fe^2+^-heme is known to bind gases. Yet several studies indicate that both Rev-Erbβ and its Drosophila analogue, E75 are involved in sensing diatomic gases.^19,22,23^

Here we demonstrate that Rev-Erbβ contains Fe^3+^-heme yet still functions as a gas sensor by coupling heme reduction to binding CO and NO. Our work explains how and why ferric hemoproteins like Rev-Erbβ are ideal gas sensors.

## Results and Discussion

### Rev-Erbβ predominantly exists in the ferric form in cells

Responsiveness of nuclear receptors to the redox status of the cell is important in matching homeostasis to the changing metabolic conditions.^16^ The thiol-disulfide and Fe^3+^/Fe^2+^-heme redox switches in Rev-Erbβ allow it to adopt variable heme coordination spheres and heme affinities.^18,20^ This dictates the protein’s heme occupancy under limiting concentrations of cellular labile heme, with Fe^3+^- and Fe^2+^-heme exhibiting K_d_ values of 0.1 and 22 nM, respectively.^9,21^

We performed spectroelectrochemical and electron paramagnetic resonance (EPR) experiments to determine the cellular resting redox state of Rev-Erbβ. By poising the protein solution at increasingly positive potentials (oxidative titrations) from −691 to 209 mV, the λ_max_ of the Soret band shifted from 427 (Fe^2+^ protein) to 422 (Fe^3+^) nm (Fig. 1b and 1c). The α bands decreased in intensity and shifted from 560 to 570 nm, with the difference spectra showing a maximum absorption decrease at 466 nm and a concomitant increase at 560 nm (extended data Fig. 1a). In some of the titrations, the λ_max_ of the Soret band for the Fe^2+^ protein was observed at 422 nm as opposed to the expected 427 nm (extended data Fig. 1b)^1^. The titrations were reversible with Nernst fits indicating the existence of two populations of the protein with fairly low redox potentials. The calculated midpoint potentials for reductive titrations were −224.9 mV and −405.0 mV and for the oxidative titrations were −229.4 mV and −363.8 mV (Fig. 1c and extended data table 1). Given that the redox potential within the nucleus is ∼ −280 mV, it appears that the protein equilibrates between the Fe^3+^ and Fe^2+^ states with the majority of heme in the Fe^3+^ state.^24^

Previously reported magnetic circular dichroism (MCD) and Resonance Raman studies of the Rev-Erbβ LBD indicated the presence of two Fe^2+^ heme populations.^18^ Our MCD experiments with longer LBD constructs (including residues 275-357) than previously reported also show the presence of a mixture of five-coordinate (5c) high-spin (HS) and six-coordinate (6c) LS Fe^2+^-heme populations. The two Fe^2+^ populations were characterized by the temperature-dependent *C*-term and the temperature-independent *A*-term respectively (extended data Fig. 2a). These MCD data agree with conclusion of the electrochemical titrations that Rev-Erbβ contains two Fe^2+^ heme populations. Our MCD experiments with the full-length and the LBD protein display identical spectral features (extended data Fig. 2b) demonstrating that the LBD retains the heme coordination sphere of the full-length protein and hence serves as a prototype for the full-length protein.^18^ Furthermore, similar MCD spectra were obtained for the protein bound to NCoR1 and to DNA (extended data Fig. 2b), indicating that the different domains of Rev-Erbβ behave as modules, rather than as tightly coupled units.^16^

**Fig. 2.**
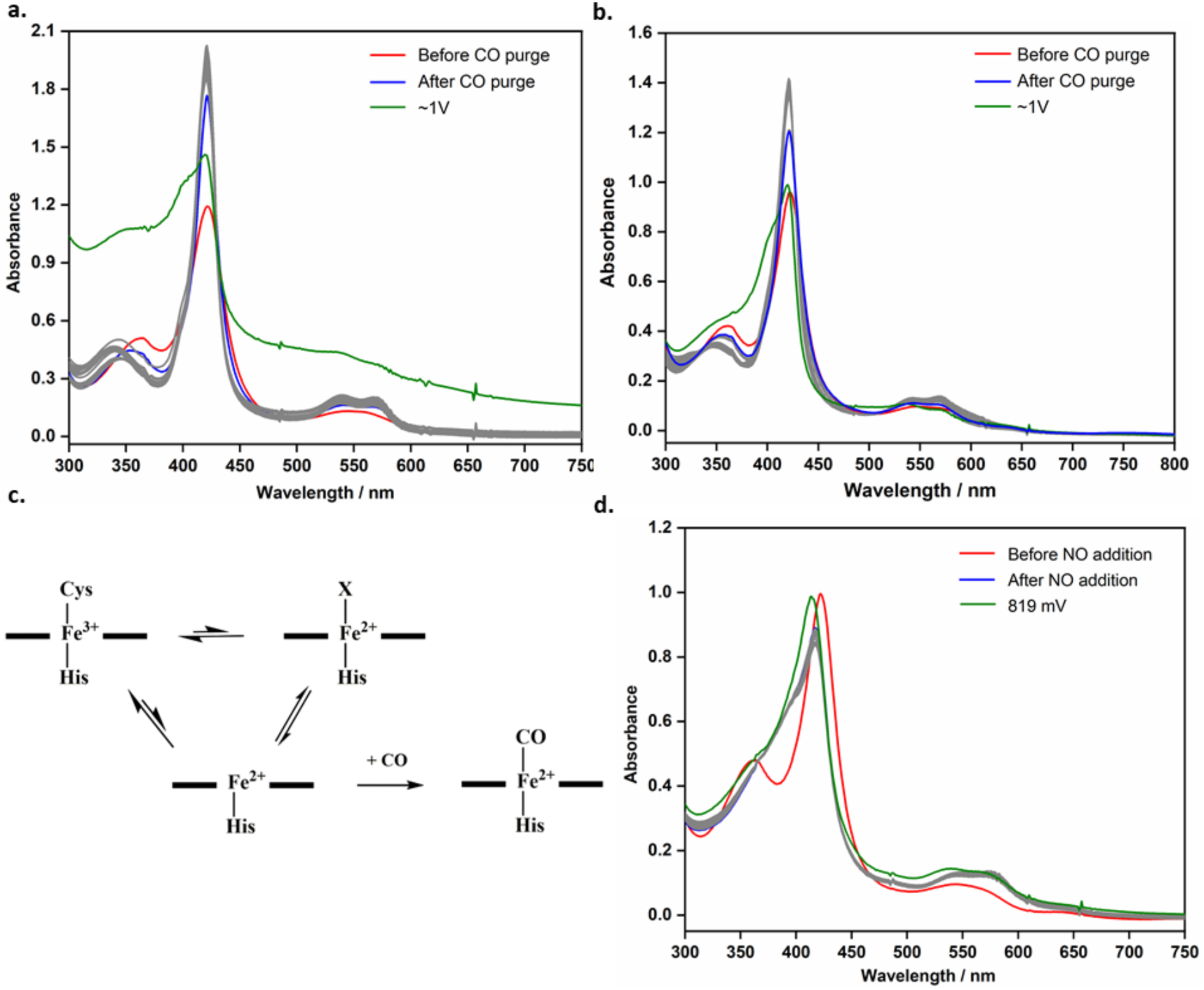
Redox Properties of Rev-Erbβ LBD in presence of CO and NO. Spectral changes observed during potentiometric titration of Rev-Erbβ LBD **a**, with low potential redox mediators under CO atmosphere, [LBD] = 118.5 μM. **b**, with high and low potential redox mediators under CO atmosphere, [LBD] = 95.5 μM. Grey traces represent absorption at potentials from 159 to 959 mV in **(a)** and −11 to 959 mV in **(b). c**, proposed scheme of CO-mediated Fe^3+^-LBD reduction. **d**, Potentiometric titration of LBD with high and low potential redox mediators in presence of NO. Grey traces represent absorption at potentials from 189 to 789 mV, [LBD] = 99.1 μM.

Given the high affinity of Rev-Erbβ for Fe^3+^ heme, in its resting state, only the Fe^3+^ form of this nuclear receptor would be heme-replete at the low nanomolar concentrations of labile heme in the nucleus.^9,21^ Our experiments suggest that Rev-Erbβ evolved its tight-binding Fe^3+^ heme binding site to accommodate to this stringent environment of heme scarcity.

Whole-cell EPR spectroscopy of *E. coli* cells over-expressing the LBD^2^ validated the spectroelectrochemical results indicating that Rev-Erbβ exists in the Fe^3+^ form under normoxic cellular redox conditions.^20^ The rhombic EPR spectrum exhibited *g*-values (g = 2.48, 2.27, and 1.88) (Fig. 1d and 1e) that are identical to those of the purified protein, consistent with His/Cys coordinated Fe^3+^-heme, as seen in the crystal structure (Fig. 1a).

We reasoned that if cellular Rev-Erbβ contained significant amounts of Fe^2+^ heme, treatment with an oxidant would enhance the EPR signal. However, addition of H_2_O_2_ increased the signal intensity of extraneous cellular heme (*g* = 4.30) but not that of the LBD Fe^3+^-heme. Instead, a modest (∼ 15 %) signal loss was observed. On the other hand, treatment with the reductant dithionite (20 mM) caused a 36 % decrease in EPR signal intensity from the LBD (Fig. 1c) as the Fe^3+^ form of the LBD was reduced to the EPR-silent Fe^2+^ form. A more pronounced (∼ 73 %) signal reduction occurred upon solid dithionite treatment for 5 min (Fig. 1d). As expected, dithionite treatment also decreased the EPR signal intensities at *g* = 4.30 and 5.96. Thus, our combined spectroscopic and electrochemical studies demonstrate that within the cell Rev-Erbβ primarily exists as 6c His/Cys ligated Fe^3+^-heme with and that binding to neither DNA or co-repressor alters this property as was apparent from the MCD experiments.

### Ferric Rev-Erbβ as CO sensor

We next focused on the apparent conundrum of how this Fe^3+^ protein could function as a gas sensor, when it is recognized that only Fe^2+^ heme can bind CO and NO with high affinity.^19,22,23^ Recognizing that the iron coordination sphere has a significant impact on its redox potential (E^0^), we reasoned that CO binding to Rev-Erbβ could increase the apparent redox potential of the Fe^3+/2+^ redox couple.^25^ CO introduction into the electrochemical setup containing low-potential dyes, resulted in a color change of the solution from orange-red (typical of the Fe^3+^ protein) to pink. The Soret absorption maximum shifted to 421 nm and the Q-bands exhibited maxima at 539 nm and 570 nm, characteristic of Fe^2+^-CO bound protein (Fig. 2a). This reaction is apparently irreversible since oxidative titrations of the CO-bound protein at potentials as high as +959 mV did not elicit any noticeable re-oxidation (Fig. 2a). We stopped the experiment at a potential of 989 mV because the protein began to precipitate. Even when both low- and high-potential redox mediators were included in the experiment (Fig. 2b), no change in the Fe^2+^-CO spectrum was observed. These results demonstrate that binding of CO increases the apparent redox potential of the Fe^3+/2+^ couple of the LBD heme by over 1.2 Volts!

The auto-reduction of heme proteins driven by a water-gas shift reaction in the absence of a reductant, has been observed in cytochrome c oxidase, hemoglobin and myoglobin.^26^ To test this possibility, we monitored conversion of ^13^CO to ^13^CO_2_ by the LBD protein using gas chromatography mass spectrometry (GCMS) and quantified the ^13^CO_2_ production in the headspace with respect to ^12^CO_2_ using CODH as a positive control.^27^ The ^13^CO_2_/^12^CO_2_ ratio for Rev-Erbβ was similar to that of control experiments lacking the LBD and was significantly lower than the CODH control (extended data Fig. 3). Thus, CO oxidation does not drive heme reduction in Rev-Erbβ.

**Fig. 3.**
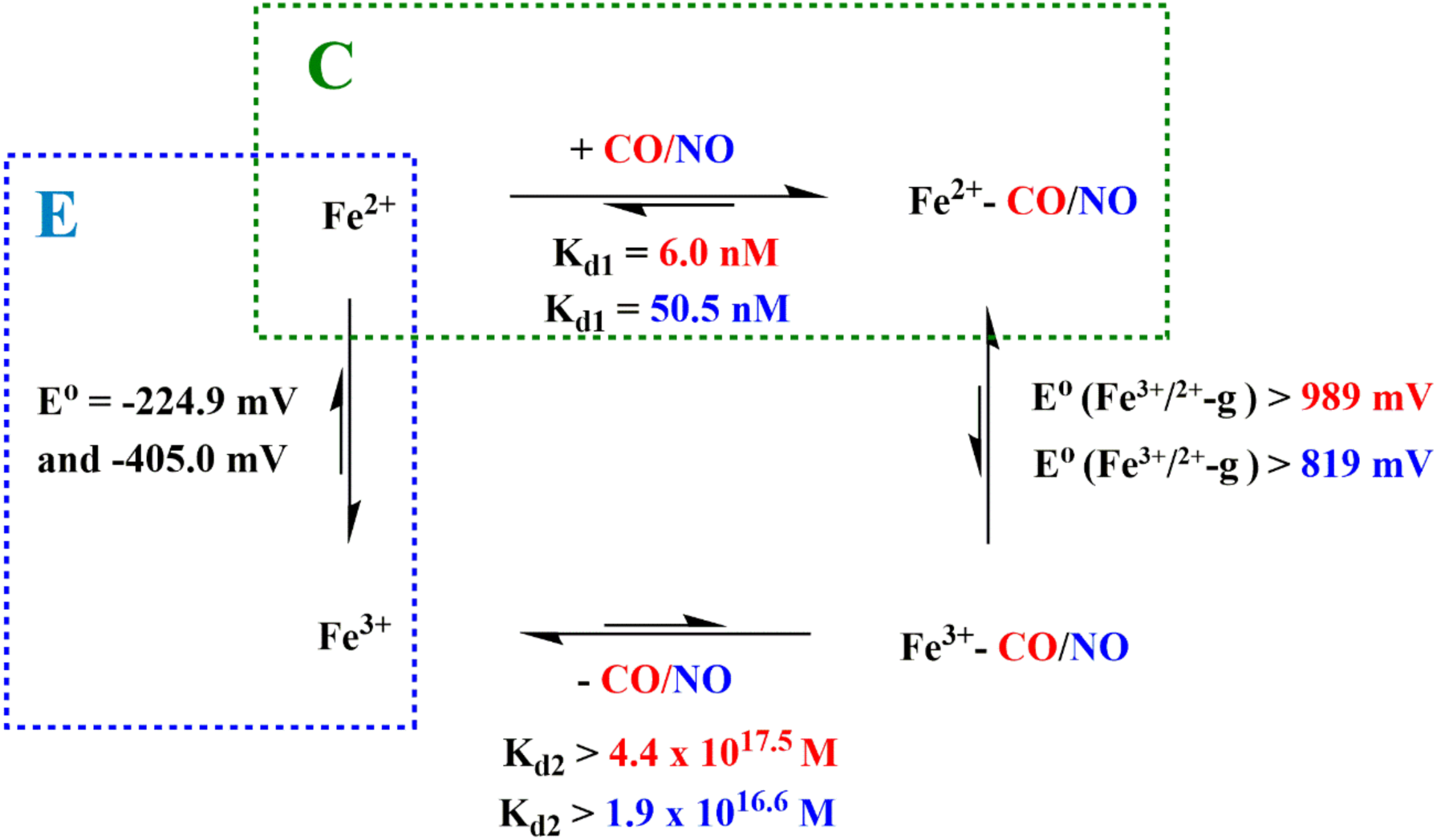
Gas driven reduction of Fe^3+^ Rev-Erbβ by EC mechanism. Thermodynamic box depicting redox potentials and gas binding constants of different redox and gas-bound states of Rev-Erbβ LBD. Red and blue indicate thermodynamic parameters for CO and NO, respectively. E^0^ and E^0^ (Fe^3+/2+^-g) represent the redox potentials of the Fe^3+^/Fe^2+^ couple in absence and presence of CO/NO respectively. K_d1_ and K_d2_ represent the binding constants of the Fe^2+^ and Fe^3+^ protein to CO/NO respectively. Coupling of electron transfer with chemical reaction is depicted by the blue and green boxes.

Formation of Rev-Erbβ Fe^2+^-CO was also not driven by the intramolecular electron transfer from Cys thiolates to Fe^3+^-heme resulting in protein dimerization under CO atmosphere, as observed with the Cys variants of myoglobin.^28^ Our SDS-PAGE analysis showed no increase in the LBD-dimer band intensity in the presence of CO (extended data Fig. 4a and 4b). Furthermore, unlike the mechanism of heme reduction delineated by Hirota S. *et*.*al*., which solely relies on protein thiolates as the electron source, the Fe^3+^ to Fe^2+^-CO transition in Rev-Erbβ requires low potential redox mediators (extended data Fig. 4c).

**Fig. 4.**
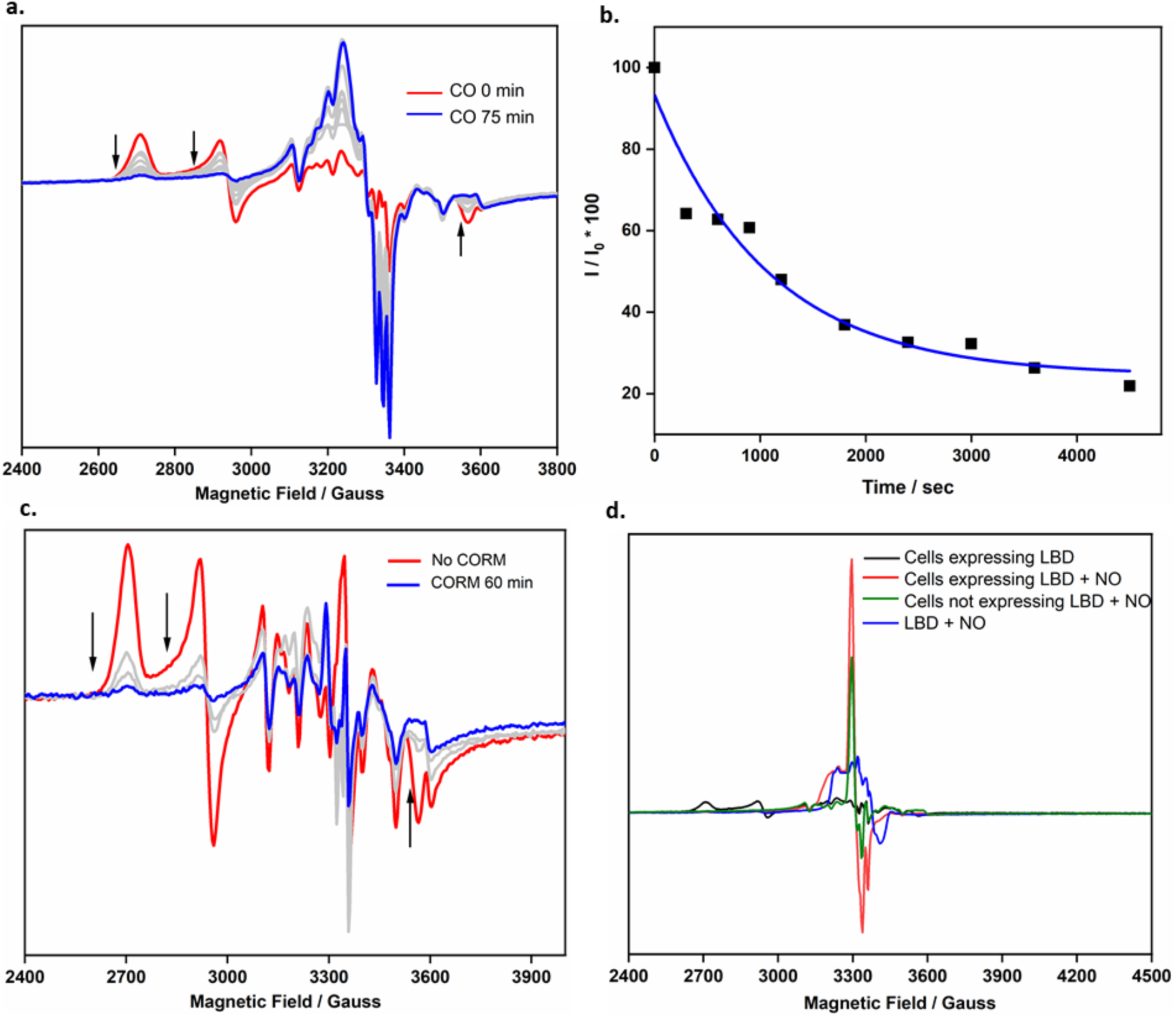
Effect of CO and NO on the redox state of Rev-Erbβ *in cellulo*. **a**, EPR spectra of *E. coli* cells over-expressing Rev-Erbβ LBD purged with CO for variable times (0, 5, 10, 15, 20, 30, 40, 50, 60 and 75 min). **b**, Scatter plots represents percentage of observed EPR signal intensity with respect to cells without CO treatment at 2708.7 G versus CO purge times. Blue line represents the fit of the data to single-exponential decay; **c**, EPR spectra of *E. coli* cells over-expressing LBD with and without 20 mM CORM-A1 treatment for variable times (20, 40, 60 min). Full EPR spectra for panels **a** and **c** are shown in extended data Fig. **7a** and 7**b**, respectively. EPR Conditions: temperature 11 K (CORM experiments) 15 K (CO purge experiments), microwave power 20 µW (CORM experiments) 208 µW (CO purge experiments), microwave frequencies of 9.3850, 9.3855, 9.3846, 9.3823, 9.3842, 9.3854, 9.3845, 9.3828, 9.3841, 9.3880 GHz (for CO purge experiments in **a**) and 9.3803, 9.3797, 9.3808, 9.3795 GHz (for CORM treatment in **c**), modulation frequency 100 kHz, modulation amplitude 7 G, 2 scans, 327.68 ms time constant; **d**, EPR spectra of (i) *E. coli* cells overexpressing LBD with and without Proline NONOate treatment (10 mM, 6 min), (ii) *E. coli* cells containing empty vector pMCSG9 after Proline NONOate treatment (10 mM, 6 min), (iii) Rev-Erbβ after Proline NONOate addition. EPR Conditions: temperature 11 K, microwave power 20 µW; microwave frequency 9.3831 and 9.3823 GHz for (i), 9.3840 GHz for (ii), 9.3849 GHz for (iii), modulation frequency 100 kHz; modulation amplitude 3 G, 4 scans, 327.68 ms time constant.

The gargantuan >1.2 V increase in midpoint potential of the Fe^3+/2+^ couple when CO binds to the LBD evokes an EC mechanism in which **E**lectrochemistry is coupled to a **C**hemical reaction (Fig. 3). At ∼ −280 mV, the redox poise within the nucleus, Rev-Erbβ exists mainly the Fe^3+^ form. The strong interaction of the residual Fe^2+^-heme with CO (K_d_ = 6 nM) exerts a driving force to push the thermodynamically unfavorable heme reduction to 100 % Fe^2+^-protein (Fig. 2c).^8^ Similarly, the high stability of the Fe^2+^-CO protein (K_d_ > 4.4 x 10^17.5^ M) excludes the formation of an Fe^3+^-CO complex (Fig. 3). To stabilize Fe^3+^-CO, Rev-Erbβ would need to be exposed to 10^40^ moles of CO, which is 24 orders of magnitude greater than the amount of CO (10^21^ liters) in the earth’s atmosphere. Regeneration of the Fe^3+^ state would not occur until CO concentrations drop below the K_d_ of 6 nM.

### Ferric Rev-Erbβ as NO sensor

When Rev-Erbβ is treated with the gaseous signaling molecule NO, its transcriptional repression activity markedly decreases, suggesting it to be an NO sensor.^19^ As with CO, NO addition to the Fe^3+^ protein resulted in formation of Fe^2+^-NO, with a blue shift in the Soret band from 422 to 417 nm (Fig. 2d). The EPR spectrum of the NO adduct showed a multi-line spectrum typical of a NO- and His-bound heme where the unpaired spin on NO couples to the coordinating His nitrogen trans to the bound-NO (Fig. 4d).^18^ Potentiometric oxidation in the presence of NO elicited no redox transition at up to 789 mV, (Fig. 2d). The protein began to precipitate at a potential of 820 mV, similar to the Fe^2+^-CO titration, indicating a much higher stability of the Fe^2+^-NO complex than the Fe^3+^-NO complex (> 10^23.6^) (Fig. 3). This high stability is imparted by the strong affinity (K_d_ = 50.5 nM) of the Fe^2+^-LBD for NO (extended data Fig. 6), which drives the thermodynamically unfavorable heme reduction. NO can reduce Rev-Erbβ in the absence of redox mediators (extended data Fig. 5a), implying that NO can possibly serve as a source of electrons *in cellulo* to facilitate the reduction of Rev-Erbβ. There are several known mechanisms of reductive nitrosylation that involve either generation of nitrite or S-NO.^29^ We do not observe formation of nitrite, suggesting an Cys-S-NO related heme reduction. Regardless, the stability conferred by NO to trap Rev-Erbβ in the reduced state may as well be a cellular mechanism to sense NO which further manifests as the biochemical signaling effects of the NO-bound protein.

Given the 220-fold higher affinity of the LBD for Fe^3+^-versus Fe^2+^-heme, having a low Fe^3+/2+^ redox potential allows Rev-Erb to be heme-replete at the low nM concentrations of labile heme in the nucleus until the cell is exposed to gaseous signaling modulators like CO and NO, when it gets reduced (extended data Fig. 8). Because of the high stability of the gas-bound Fe^2+^ state, recycling Rev-Erb to its resting Fe^3+^ state would occur when the cellular concentrations of CO or NO drop below the K_d_-value. Thus, Rev-Erbβ (and potentially other heme-based signaling proteins) seems ideally poised for its function as a heme- and gas-regulated nuclear receptor to control such myriad processes as regulation of the circadian rhythm, glucose and lipid metabolism, and inflammatory responses.

### Rev-Erbβ Fe^3+^-Heme: A gas sensor in cells

To complement the *in vitro* experiments, we used *in vivo* EPR to determine if CO and NO convert Fe^3+^-heme to the EPR-silent Fe^2+^-CO or Fe^2+^-NO state *in cellulo*. After purging with CO, *E. coli* cells over-expressing LBD exhibited time-dependent loss of the Fe^3+^-heme EPR signal (Fig 4a). A 36% reduction in signal intensity of the characteristic low-spin Fe^3+^-heme occurred within 5 mins of CO purge and followed a single exponential decay with rate constant of 9.3 x 10^−4^ s^-1^ (Fig. 4a and 4b). Surprisingly, CO purging also resulted in an EPR spectral feature with *g-* and triplet hyperfine splitting values representative of a 5c Fe^2+^-Heme-NO complex.^30^ The concurrent decrease in HS-heme, as shown by the loss of the *g*= 6 EPR signal intensity (extended data Fig. 7c), can possibly account for formation of the NO-bound heme. CO and NO are recognized gasotransmitters that likely intersect as regulators of bacterial metabolism; however, this observed CO-mediated NO release needs further investigation.^31^

When cells were treated with the CO-releasing molecule CORM-A1, the Fe^3+^-heme EPR signal decreased by 70% within 20 mins (Fig. 3c, extended data Fig. 7e) and nearly disappeared (93.0 %) after 60 min of CORM treatment. CORM treatment also decreased the 6-line spectrum cellular Mn^2+^ and high-spin heme, and modestly reduced the signal of extraneous non-heme iron (g = 4.3) (extended data Fig. 7f, 7g).

Addition of proline NONOate, which generates NO, to LBD-overexpressing *E. coli* cells elicits a response similar that of CO. NO treatment results in a concentration-dependent loss of the Rev-Erb Fe^3+^-heme signal (extended data Fig. 5b). This reduction is accompanied by gain of a signal at *g* = 2.06 and 2.03 (Fig. 4d). The signal increase at *g* = 2.03 is also observed for control cells containing the empty vector (extended data Fig. 5c) and can be attributed to the reaction of NO with iron-sulfur clusters yielding dinitrosyl iron complexes.^32^ The signal build-up at *g* = 2.06 is also observed upon the treatment of the LBD with NO (Fig. 4d), confirming nitrosylation of the Fe^3+^-heme of Rev-Erbβ *in cellulo*.

## Conclusions

Our *in vitro* and whole-cell experiments demonstrate that the low Fe^3+/2+^-heme redox potential of Rev-Erbβ dictates that it rests in the Fe^3+^ state in cells allowing the protein to be heme-replete. Our work also establishes how Fe^3+-^Rev-Erbβ functions as a gas sensor when signaling gases (CO, NO, O_2_) selectively bind the Fe^2+^ state of metallocofactors.^17^ Rev-Erbβ takes advantage of this high selectivity by linking heme reduction (**e**lectrochemistry) to gas binding (**c**hemistry) via an EC mechanism. This selectivity manifests as an increase in the apparent redox potential of the Fe^3+^/Fe^2+^ couple in presence of the gases. Our findings also explain how Rev-Erbβ, which rests in the Fe^3+^ state, is de-repressed by NO in mammalian cells.^19^

The results described here impact a relatively ignored aspect of metal ion homeostasis in which the resting tight-binding oxidized state of a metallocofactor is recruited into its active state by the strategy of chemical coupling. Coupled reactions drive several fundamental biological processes where redox changes are accompanied by ligand, substrate and proton binding and/or conformational changes.^33-35^ Not only signaling gases but various substrates or ligands can be interfaced to the Fe^3+/2+^ redox couple. A well-studied example is Cytochrome P450, which because of its low Fe^3+/2+^ potential is predominantly in the Fe^3+^ state within the cell.^36^ Substrate binding increases this potential, promoting reduction to the Fe^2+^ state, which binds O_2_ and initiates the formation of reactive iron-oxo catalytic intermediates involved in substrate oxidation. Our findings offer a fundamental explanation as to how and why chemical gating of electron transfer in hemoproteins like Rev-Erbβ allows them to exist in the oxidized form but still function as a gas-sensor.^37-40^ Often, a conformational change accompanies reduction and/or ligand binding. CO and NO are increasingly recognized as important regulators of the circadian rhythm. However, the molecular mechanisms underlying the same remain elusive. We anticipate that this work will pave the way for investigating how CO/NO can be integrated into the circadian rhythm framework due to their ability to cause redox reactions triggering conformational changes in metalloproteins like Rev-Erbβ.

## Materials and Methods

Unless otherwise mentioned, all chemicals were of analytical grade, obtained from commercial sources, and used without further purification. All experiments were performed in 0.5x TNG (25 mM Tris-HCl, 150 mM NaCl and 5% glycerol) pH 8.0 buffer unless otherwise mentioned. Experiments involving the use of Proline NONOate and CORM-A1 were performed in 0.5x TNG pH 7.4 buffer. All absorption measurements were carried out on an Agilent Cary 8454 UV-visible spectrophotometer inside the anaerobic chamber with less than 0.3 ppm dioxygen except for Bradford assays and heme concentration measurements, which were performed aerobically on a Shimadzu UV-2600 spectrophotometer. The Rev-DR2 DNA sequence was obtained from Integrated DNA Technologies (Coralville, IA).

### Protein expression and purification

The Rev-Erbβ ligand binding domain (LBD) encompassing residues 370-579 was expressed as described previously.^9^ Briefly, the Rev-Erbβ LBD is expressed as a translational fusion between maltose binding protein (MBP) and the LBD protein separated by an internal tobacco etch virus (TEV) protease cleavage site. The fusion protein was expressed in Escherichia coli BL21(DE3) cells grown at 37 °C in Terrific Broth supplemented with 0.4% glycerol and 200 µg ml^-1^ ampicillin. Protein expression was induced with 0.5 mM isopropyl β-D-1-thiogalactopyranoside (IPTG). The cell pellet obtained by centrifugation was suspended in TNG buffer (50 mM Tris-HCl, pH 8.0, 300 mM NaCl, and 10% glycerol) containing 1 mM dithiothreitol, 6 mM benzamidine, 1 mM EDTA, 0.5 mM phenylmethanesulfonyl fluoride, and 1x protease inhibitor mixture (Roche Applied Science) on ice and lysed by sonication. The cell-free extract obtained post centrifugation was applied to an amylose column (New England Biolabs) equilibrated in TNG with 1 mM DTT at 4 °C. After washing the column with the TNG buffer, the pure fusion protein was eluted with TNG containing 20 mM maltose. MBP tag was removed by TEV protease treatment. Removal of the cleaved MBP tag was achieved by passing the protein solution through a hydroxyapatite column. The protein was eluted using a buffer containing 20 mM sodium phosphate, 200 mM NaCl, pH 7.2. The eluted protein was concentrated and buffer exchanged into 0.5 x TNG buffer using an Amicon ultrafiltration device (Millipore) containing a 10 kDa membrane.

The heme content in the protein was determined by the pyridine hemochrome assay.^41^ The heme content varied from 10 to 20 % approximately between different batches.

MCD experiments were performed with slightly longer LBD constructs (residues 247-57) of Rev-Erbβ than previously reported. The LBD was expressed and purified as an N-terminal His_6_-tagged protein according to the previously reported protocol.^20^ MBP-NCoR1 protein containing all three interaction domains and full-length MGC Rev-Erbβ used for MCD studies were purified according to the previously reported protocol.^8^

### Preparation of thiol-reduced protein

Thiol-reduced protein was prepared in an OMNI-LAB anaerobic chamber (Vacuum atmosphere) maintained with N_2_ atmosphere containing < 0.3 ppm dioxygen. The protein was treated with > 10-fold excess of TCEP in DMSO (Sigma) and incubated for an hour in the anaerobic chamber at room temperature. The thiol-reduced protein was exchanged into anaerobic 0.5x TNG buffer using Microbiospin 6 columns (Bio-Rad). The protein concentration was determined by the Bradford assay (ThermoFisher scientific). To prepare the heme-bound protein, a primary heme stock was prepared by dissolving hemin in 0.1 M NaOH and 10% DMSO. The solution was filtered through 0.2 μm DMSO-safe synrige filters (Pall Corporation) and diluted to ∼ 1 mM in 0.5x TNG. The concentration of the secondary stock was determined from the absorption at 385 nm (ε = 58.44 mM^-1^ cm^-1^).^42^ Since the heme content of the protein varied from 10 to 20 % between batches the thiol reduced protein and external heme were mixed in the ratio of 1 : 0.6 or 1 : 0.5, respectively under anaerobic conditions. For all experiments, heme was added at sub-stoichiometric concentration to the protein to avoid any spurious effect from unbound heme. Concentrations of protein reported are that of the heme bound protein.

### Redox titrations

Redox titrations were performed at 25 °C in an anaerobic chamber with < 0.3 ppm O_2_. All experiments were performed in 0.5x TNG anaerobic buffer. Two cocktails of mediator dyes (with 1 mM of each dye)^3^, low potential dyes and high potential dyes, were prepared in 0.5x TNG buffer (pH 7.4 or 8.0) for the redox titrations. Triquat and dimethyl triquat were synthesized according to previously reported protocol.^43^

Low Potential Dyes
| Dyes | Potential* | Reference |
| --- | --- | --- |
| Potassium Indigotetrasulfonate | -46 mV | 44 |
| Indigo Carmine | -125 mV | 44 |
| Anthraquinone-1,5-disulfonic acid | -200 mV | 45 |
| Anthraquinone-2-sulfonic acid | -225mV | 39 |
| Benzyl Viologen | -358 mV | 39 |
| Methyl Viologen | -440 mV | 39 |
| Triquat | -549 mV | 46 |
| Dimethyl triquat | -684 mV | 46 |

High Potential Dyes
| Dyes | Potential* | Reference |
| --- | --- | --- |
| Methylene Blue | 11 mV | 44 |
| Methyl Hydroquinone | 460 mV | 47 |
| Thionine | 56 mV | 44 |
| N-Acetyl L-tyrosinamide | 650 mV | 47 |
| Adenine | 750 mV | 47 |
| Potassium Indigotetrasulfonate | -46 mV | 44 |
| Indigo Carmine | -125 mV | 44 |
| Methylene Green | 123 mV | 48 |
| Bindscheder's Green | 224 mV | 44 |

\*All potentials are reported versus the standard hydrogen electrode (SHE).

The redox mediator dyes were added to the thiol-reduced protein at molar ratios of 1:10. The redox titrations were carried out in an electrochemical cell (BASI) equipped with three electrodes (Au gauze as working electrode, Pt as auxillary electrode, Ag/AgCl as reference electrode) connected to a CV-27 voltammogram (BASI). The applied potential was varied from 0 to −900 mV vs Ag/AgCl. The redox titrations were followed by monitoring the absorption changes of the protein inside the anaerobic chamber. The absorption spectrum reported at each potential was recorded after no further change in absorption was observed and/or the current reached 0 µA. The LBD heme redox potential (E°) was determined from the Nernst-plot of the fraction of protein reduced versus the applied potential. The data best fit to a one-electron redox reaction with two populations (equation S1), where [red] and [oxd] represent concentrations of reduced and oxidized protein, respectively; E°_1_ and E°_2_ represent the redox potential of the two populations; and a and b represent the amplitude of two populations with the constraint, a + b = 1. All the redox potentials are reported against SHE unless otherwise mentioned.

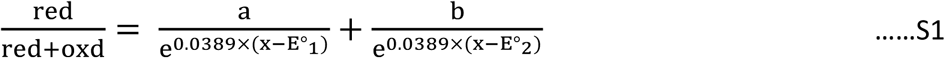

Redox titrations in presence of CO where done by purging the headspace of the electrochemical cell with CO. CO was also purged intermittently to maintain a positive pressure of CO throughout the experiment. Proline NONOate (Cayman Chemicals) was used as the source of NO for redox titrations. Proline NONOate stock solutions were prepared in 10 mM NaOH. The applied potential was varied from 209 to −691 mV for the reductive and oxidative titrations in absence of gases. Redox titrations in presence of CO were carried out in the presence of low potential dyes and a combination of low and high potential dyes with the applied potentials for the two experiments varying from 159 to 1 V and from −11 to 989 mV, respectively. Redox titrations in presence of excess NO (∼ 2 mM) were carried out in the presence of both low and high potential dyes with the applied potential varying from 189 to 819 mV.

### MCD studies

All MCD experiment samples were prepared in 25 mM Tris, 150 mM NaCl, 50 % glycerol (added as a glassing agent), pH 8.0 buffer. The LBD ferrous protein preparations also had 5 mM dithionite in the solution. The proteins of interest were injected between two quartz plate windows housed in a custom-made MCD sample holder. The samples were frozen in liquid nitrogen to produce glass.

An OXFORD SM4000 cryostat and a JASCO J-815-CD spectrometer were used for the MCD setup. The SM4000 cryostat consisting of a liquid helium-cooled superconducting magnet provided horizontal magnetic fields of 0-7 T. The J-815 spectrometer uses a gaseous nitrogen-cooled xenon lamp and a detector system consisting of two interchangeable photomultiplier tubes in the UV-vis and NIR range. The samples were loaded into a 1.5-300 K variable temperature insert (VTI), which offers optical access to the sample via four optical windows made from Suprasil B quartz. The MCD spectra were measured in [θ] = mdeg and manually converted to Δε (M^-1^ cm^-1^ T^-1^) using the conversion factor

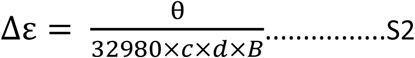

where *c* is the concentration, *B* is the magnetic field, and *d* is the path length. The product *c*x*d* can be substituted by A_MCD_/ε_UV-vis_, where A_MCD_ is the absorbance of the sample measured by the CD spectrometer and ε_UV-vis_ is the molar extinction coefficient. Complete spectra were recorded at indicated temperatures and magnetic fields.

### SDS PAGE Analysis

SDS PAGE analysis to probe protein dimer formation under CO atmosphere was performed on a gradient 4-20% SDS gel in the absence of any thiol reducing agent. Equal concentration and volumes of thiol reduced LBD protein were incubated at room temperature under N_2_ or CO atmosphere for 21 h before running gel electrophoresis and measuring the absorption spectrum of the sample. The gel images were obtained on the ChemiDoc MP Imaging system (Biorad) and the intensity analysis of the bands was done using FijiJ.

### GC-MS studies

^13^CO_2_ and ^12^CO_2_ detection were performed on an Agilent 7980B Gas Chromatograph coupled to 5977B Mass Spectral Detector. The thiol-reduced protein (60 μM) was prepared as described previously. CODH (4.8 μM) was used as a positive control. The samples (600 μl) were prepared and transferred in the anaerobic chamber in vials septum-sealed vials that were then crimp sealed with an aluminum cover (Agilent) and degassed before the introduction of CO. CO was bubbled into the solution for 105 s. 500 μl of the sample was injected and passed through a Carboxen-1010 PLOT Capillary GC column (Sigma, 30 m x 0.32 mm) kept at 25°C. Helium was used as the carrier gas. The mass spectrometer was used in selected ion monitoring (SIM) mode for detection of ^12^CO_2_ and ^13^CO_2_ at *m/z* of 44 and 45, respectively. The samples were run for a total time of 10.8 min with data acquisition from 5 to 10.8 min. The retention time for CO_2_ was 9.9 min.

The extent of conversion of ^13^CO to ^13^CO_2_ was reported as the ratio of the area under the peaks. The percentages of ^13^CO_2_ with respect to ^12^CO_2_ represent the average ± S.D. from 3 to 4 acquisitions.

### NO binding assay

The binding affinity of Fe^2+^-LBD protein for NO was determined by UV-visible spectrophotometry. The spectrophotometric studies were carried out on an Agilent Cary 8454 UV-vis diode array spectrophotometer in a quartz cuvette having a path length of 1 cm at room temperature. The experiments were performed in the anaerobic chamber in 0.5x TNG buffer, pH 7.4. Proline NONOate stocks prepared in 10 mM NaOH was used as the source of NO. Thiol-reduced protein (∼0.3 mM, 60 μl) was treated with 1 M dithionite (5.5 μl) for 30 min to reduce the ferric protein to Fe^2+^. Attempts to carry out titrations in the presence of dithionite resulted in severe precipitation and the requirement for unreasonably high amounts of NO to reach saturation indicating an interaction between dithionite and NO. Hence, dithionite was removed from the protein by two consecutive buffer exchange using Microbiospin6 columns. The buffer exchanged-protein was diluted to 3 ml for spectrophotometric titrations. Increasing amounts of NO were titrated to the protein solution and the absorption spectrum of the protein was monitored until the equilibrium was reached. The Proline NONOate stock concentration was determined by the absorption at 252 nm (ε = 8400 M^-1^ cm^-1^). The absorption of the Soret band maximum was plotted as a function of the concentration of NO (assuming 1 molecule of proline NONOate releases 2 molecules of NO) and the data obtained were fit to equation S3.

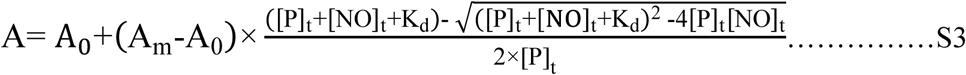

where A is the observed absorption of the protein-NO complex, A_0_ is the absorption of the protein in absence of any NO, A_m_ is the absorption of the completely NO bound protein, [NO]_t_ is the total NO concentration, and [P]_t_ is the protein concentration. [P]_t_ in Equation S3 was left as a variable, as the inflection point of the fit (representative of [P]) was sometimes lower than the known concentration of protein in the cuvette. K_d_ is reported as mean ± s.d. from 5 experiments.

### CO concentration determination

CO concentration of the buffers/solutions was determined by titration against reduced equine Myoglobin (Sigma). Experiments were carried out in a sealed cuvette with minimum headspace for any gas exchange. The CO concentrations were estimated from the percentage conversion of ferrous Myoglobin to ferrous CO Myoglobin assuming stoichiometric binding to Myoglobin. The concentrations of Fe^2+-^Myoglobin and Fe^2+^-CO Myoglobin were determined from the absorption at 435 nm (ε = 121 mM^-1^ cm^-1^) and 424 nm (ε = 207 mM^-1^ cm^-1^), respectively.

### EPR spectroscopy

EPR spectra were recorded at 11 and 15 K on an X-band Bruker EMX spectrometer (Bruker Biospin Corp.) containing an Oxford ITC4 temperature controller. All EPR samples were prepared in 0.5x TNG buffer. The samples were frozen in liquid nitrogen before experiments. Care was taken to ensure the use of the same internal and external diameter quartz tubes for EPR experiments as well as the same volumes of the sample for quantification purposes.

#### EPR of the protein

EPR of the thiol-reduced protein was carried out to test the integrity of the protein as well as quantification purposes. Thiol reduced protein was prepared as described previously. The EPR sample for the protein was prepared in the anaerobic chamber with < 0.3 ppm dioxygen. The heme content of the protein was determined using the pyridine hemochrome assay.

#### Whole-cell EPR

Whole-cell EPR was carried out for the detection/quantification of the Fe^3+^-LBD complex *in vivo. E. coli* cells expressing the LBD were inoculated from a starter culture into a secondary culture. The secondary culture was grown in the presence of 25 µM heme (heme stocks were prepared in DMSO as described previously). The cells were harvested by centrifugation. The pellet obtained from a 1 L culture was re-suspended in ∼ 5 ml 0.5x TNG buffer, pH 7.4 or pH 8.0 depending on the experiment. The cell suspension was diluted two folds including treatments with all the chemicals, for the final EPR experiments. Control experiments were carried out with *E. coli* cells containing empty vector pMCSG9 without any external supplementation of heme in the culture. CORM-A1 (Sigma) and proline NONOate (Cayman chemicals) were used as sources of CO and NO respectively. CORM-A1 is a metal ion-independent CO releasing molecule with a half-life of 21 mins at 37° C and pH 7.4. The chemical concentrations and times of incubation are indicated in the figure legend of Fig. 4. Acquisition parameters for individual samples/sets of experiments have been mentioned in the respective figure legend. CORM treatment also resulted in a decrease in the EPR signal intensity of the cellular Mn^2+^ complex (6-line spectra between 2970 G and 3655 G) (extended data Fig. 7f). Therefore, the ratio of the EPR intensity of the Fe^3+^ LBD normalized to cellular Mn^2+^ complexes was also calculated and depicted in extended data Fig. 7d.

Stocks of dithionite (1 M) and H_2_O_2_ (1 M) were prepared in 1 M Tris pH 8.0 and water, respectively. Experiments involving exogenous administration of CO gas to *E. coli* cells were carried out in sealed serum vials (Agilent) maintained at a temperature of 37 °C. The samples were continuously purged with CO gas. Equal volumes of samples were withdrawn at different periods and immediately frozen in EPR tubes. Treatment of *E. coli* cells with CORM A1 and proline NONOate were also carried pout in sealed serum vials maintained at a temperature of 37 °C. Primary stocks of CORM-A1 and proline NONOate were prepared in 10 mM NaOH. Cell-samples were withdrawn from sealed vials at different time points for the time-dependent studies of CORM treatment on *E. coli* cells.

#### Quantification of LBD-bound heme in whole cells

The percentage increase or decrease in the EPR signal observed in dithionite and H_2_O_2_ treated cells has been reported based on the EPR signal intensity change at *g*z (2.49) of the rhombic spectrum exhibited by the protein Fe^3+^-heme.

### Data availability

The derivation of some of the thermodynamic parameters in Fig 3 is included in the supplementary information. All other data supporting the findings in this study are available from the corresponding author upon request.

## Acknowledgments

CODH protein was kindly donated by Daniel Esckilsen, Ragsdale laboratory, University of Michigan. The authors acknowledge the technical help received from Thomas Yavaraski, Department of Civil and Environmental Engineering, University of Michigan, for the GCMS experiments. The funding for the research was supported by the National Institute of Health Grant R01-GM123513 (to S.W.R.).

## Author contributions

S.W.R., A.S., and E.C. conceived the project. A.S., E.C., J.B.H., and A.C.S. performed the experiments. J.B.H., A.C.S., and N.L. helped in the acquisition and analysis of the MCD data. A.S. and S.W.R. wrote the manuscript with help from all the co-authors.

## Additional information

Supplementary information is available for this paper at https://doi.org/……

## Correspondence and requests for materials

should be addressed to S.W.R.

## Extended Data Figures

**Extended Data Fig. 1.**
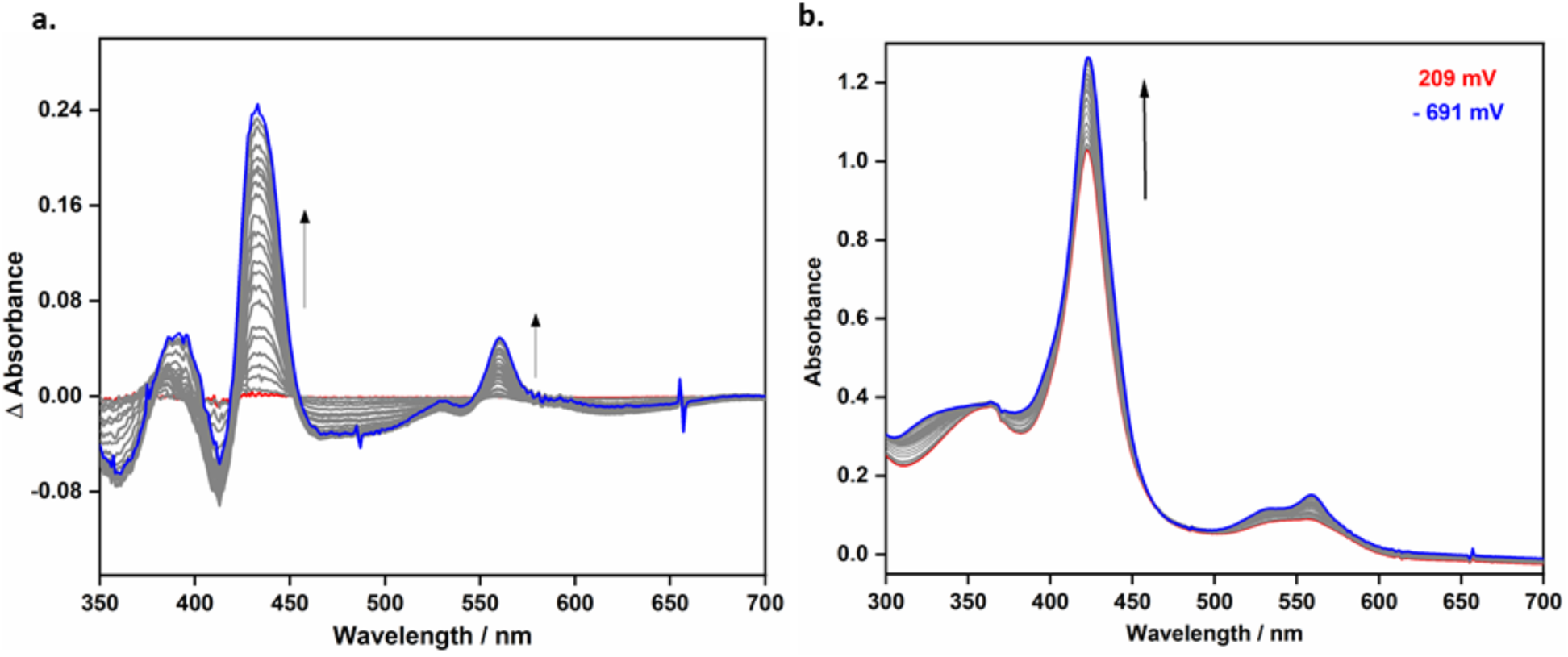
Redox Properties of Rev-Erbβ LBD. **a**, Difference spectra of the data represented in Fig. 1b, calculated by subtracting the absorption at 209 mV from absorption spectra at different potential, with the difference spectra at either the most reduced (−691 mV) and oxidized (+109 mV) represented as blue and red traces, respectively. The difference spectrum at intermediate potentials of 9, −91, −121, −141, - 181, −211, −241, −261, −281, −301, −321, −341, −361, −381, −391, −411, −431, −461, −501, −581, −641 mV are depicted in black. Arrows indicate the direction of increasing ΔA with decreasing applied potential; **b**, Spectral change observed in some of the potentiometric titrations where the Soret band did not show a shift in λ_max_. The red and blue traces are the spectra collected at the extremes of 209 and −691 mV, and the black traces at intermediate potentials.

**Extended Data Table 1.** I Redox Properties of Rev-Erbβ LBD. Comparison of the mid-point potentials of the reductive and oxidative titrations. Data from Fig. 1b was fit using equation S1.

|  | Reductive Titration | Oxidative Titration |
| --- | --- | --- |
| Amplitude of low redox potential population | $0.41 \pm 0.03$ | $0.29 \pm 0.04$ |
| Low Redox Potential (mV) | $-224.9 \pm 9.8$ | $-229.4 \pm 14.7$ |
| Amplitude of high redox potential population | $0.59 \pm 0.03$ | $0.71 \pm 0.04$ |
| High Redox Potential (mV) | $-405.0 \pm 6.7$ | $-363.8 \pm 5.7$ |

**Extended Data Fig. 2.**
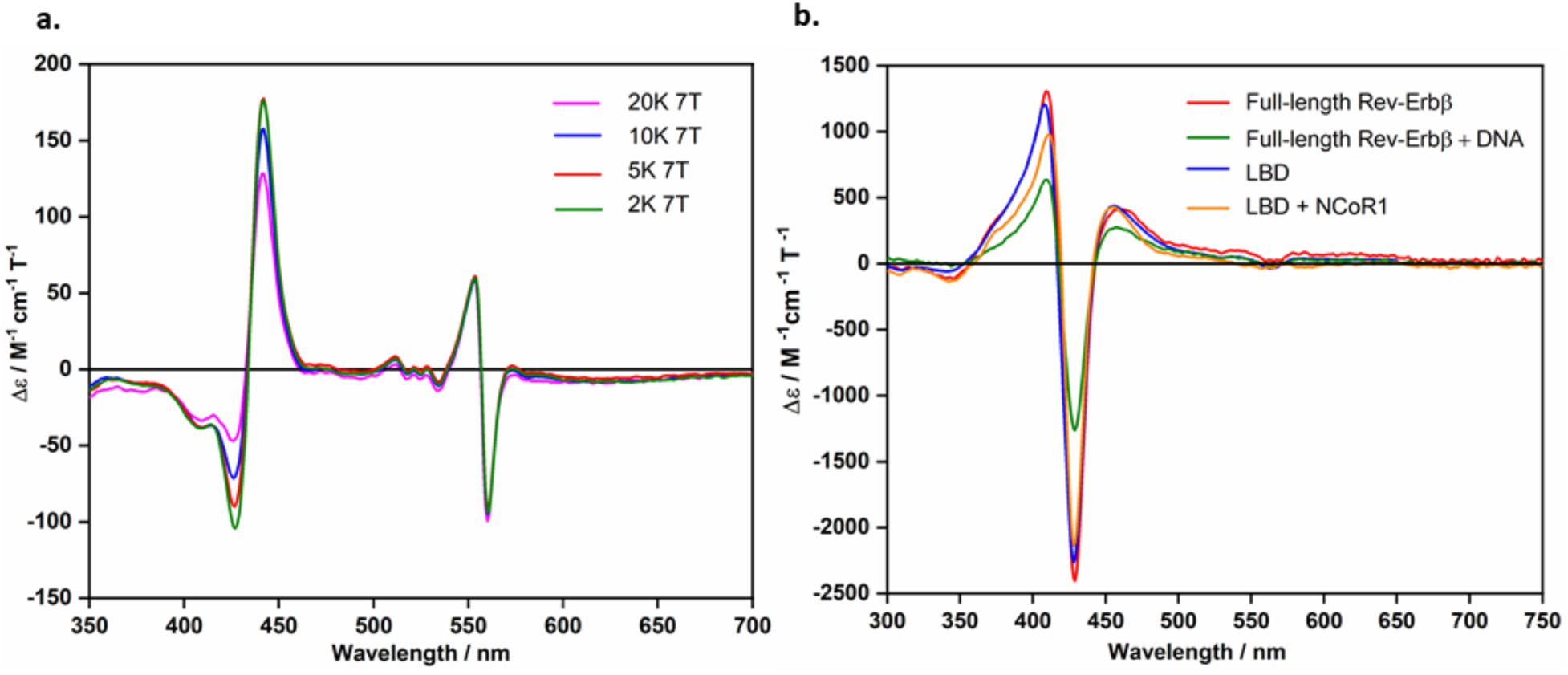
Fe^2+^ LBD exists as the mixture of 5c and 6c heme and Fe^3+^ LBD retains the heme coordination sphere of the full-length protein and full-length protein bound to DNA and LBD bound to Co-repressor. MCD spectra of **a**, Fe^2+^ LBD (27.5 μM) at different temperature and constant magnetic field (7 T); **b**, Fe^3+^ LBD (blue, 29.7 μM), Fe^3+^ Full-length protein (red, 40.3 μM), Fe^3+^ LBD (20.9 μM) with NCOR1 (20.9 μM), Fe^3+^ full-length protein (33.3 μM) with Rev-DR2 (33.3 μM) DNA^8^ sequence at 2 K; Magnetic field 1 T.

**Extended Data Fig. 3.**
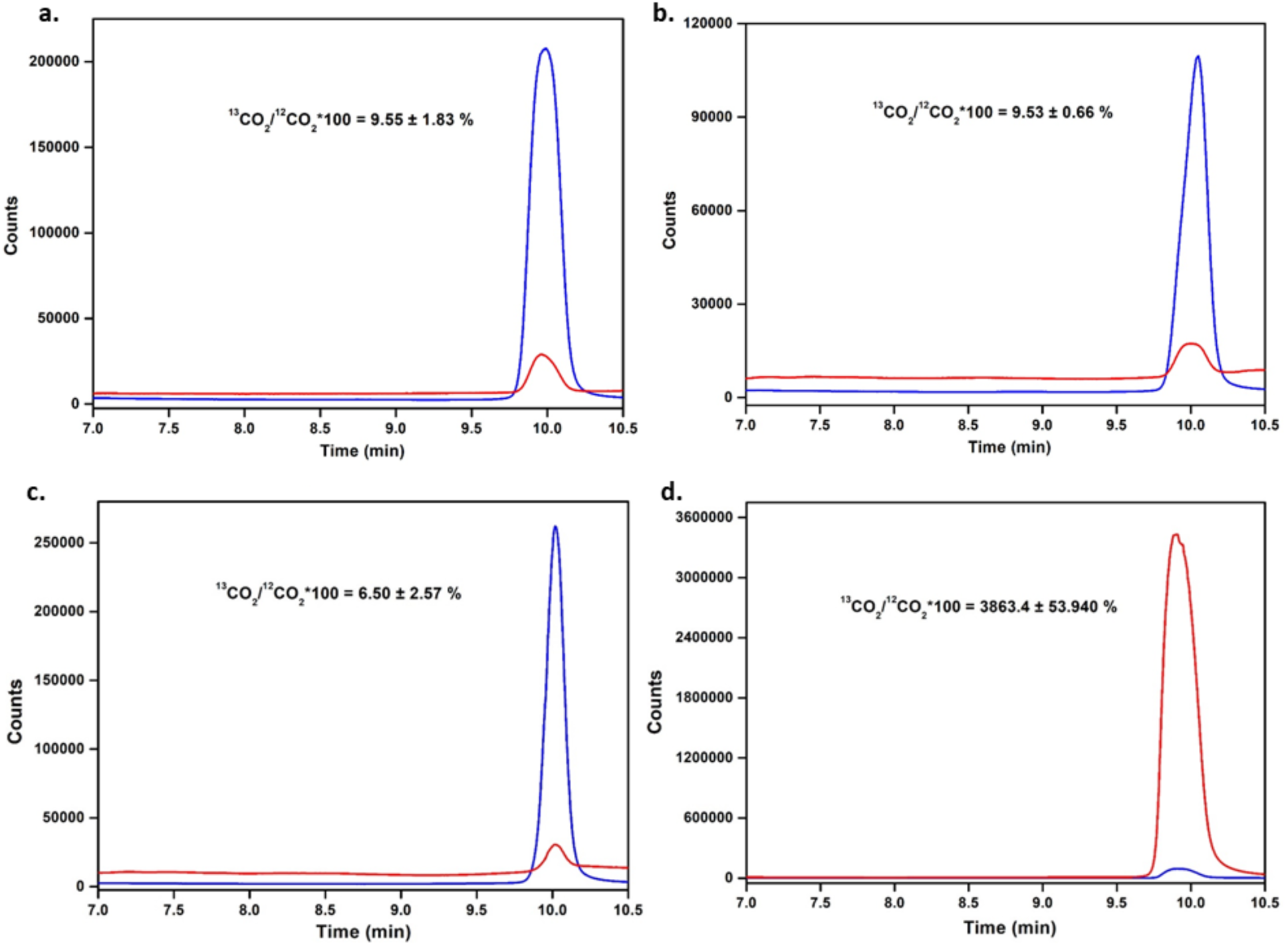
CO oxidation to CO_2_ does not drive Rev-Erbβ LBD reduction. GCMS chromatogram of gaseous ^13^CO_2_ (red) and ^12^CO_2_ (blue) in the headspace of vials containing **a**, Labelled CO; **b**, Buffer containing 10 µM low potential redox mediator dyes; **c**, Rev-Erbβ (100 μM) and low potential redox mediator dyes (10 μM); **d**, CODH (4.8 µM), low potential redox mediator dyes (10 µM), dithiothreitol (1 mM).

**Extended Data Fig. 4.**
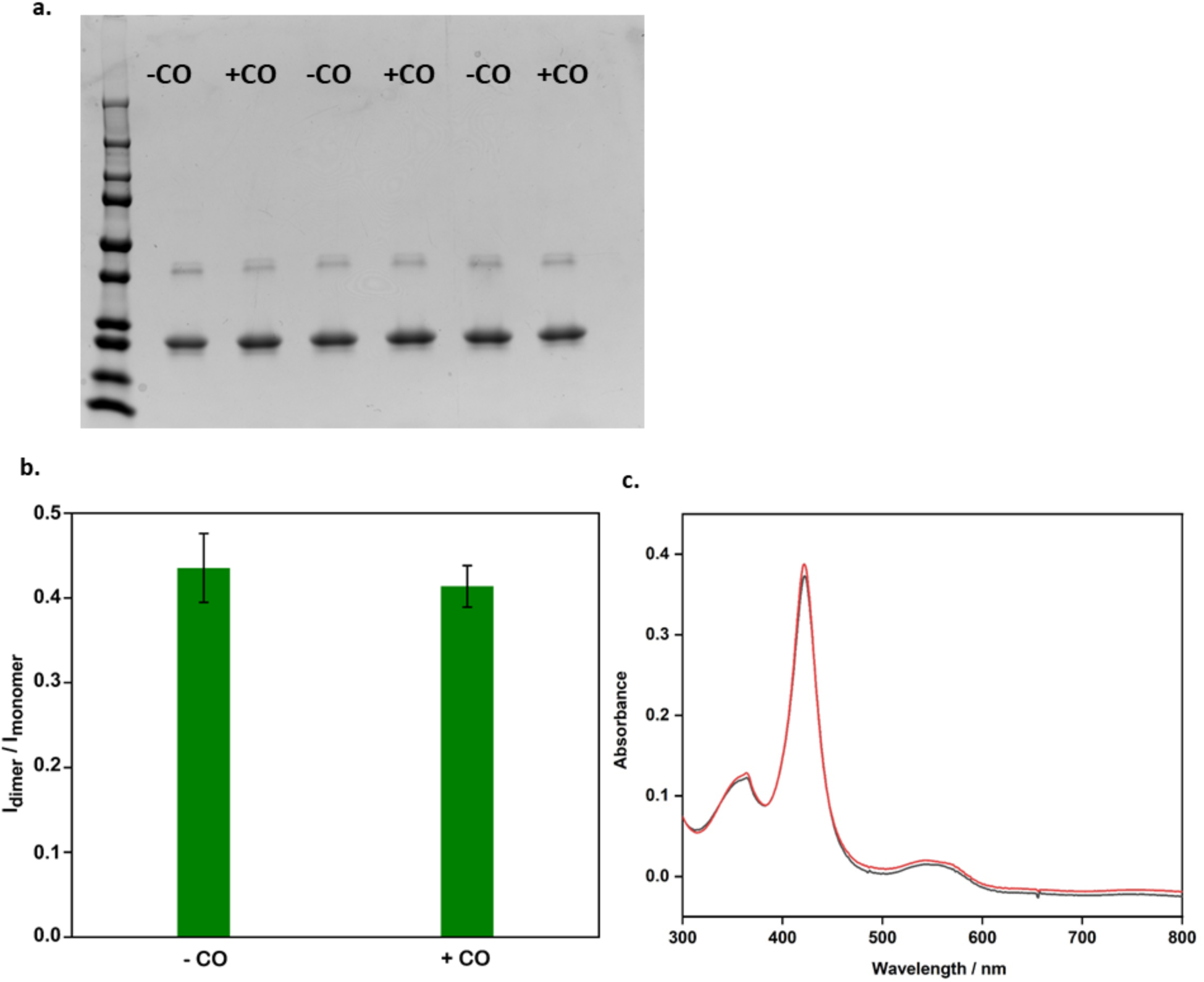
Intramolecular electron transfer from thiolates does not drive Rev-Erbβ LBD reduction. **a**, SDS-PAGE analysis of Rev-Erbβ incubated under anaerobic (-CO) and CO (+CO) atmosphere at 37 °C for 21 h, L indicates protein ladder, SDS gel analyses were repeated in triplicate; **b**, Bar plots representing the ratio of average intensity of the dimer versus monomer protein band in absence and presence of CO; **c**, Absorption spectrum of Rev-Erbβ LBD in presence and absence of CO, [Rev-Erbβ] = 37 µM without redox mediator dyes.

**Extended Data Fig. 5.**
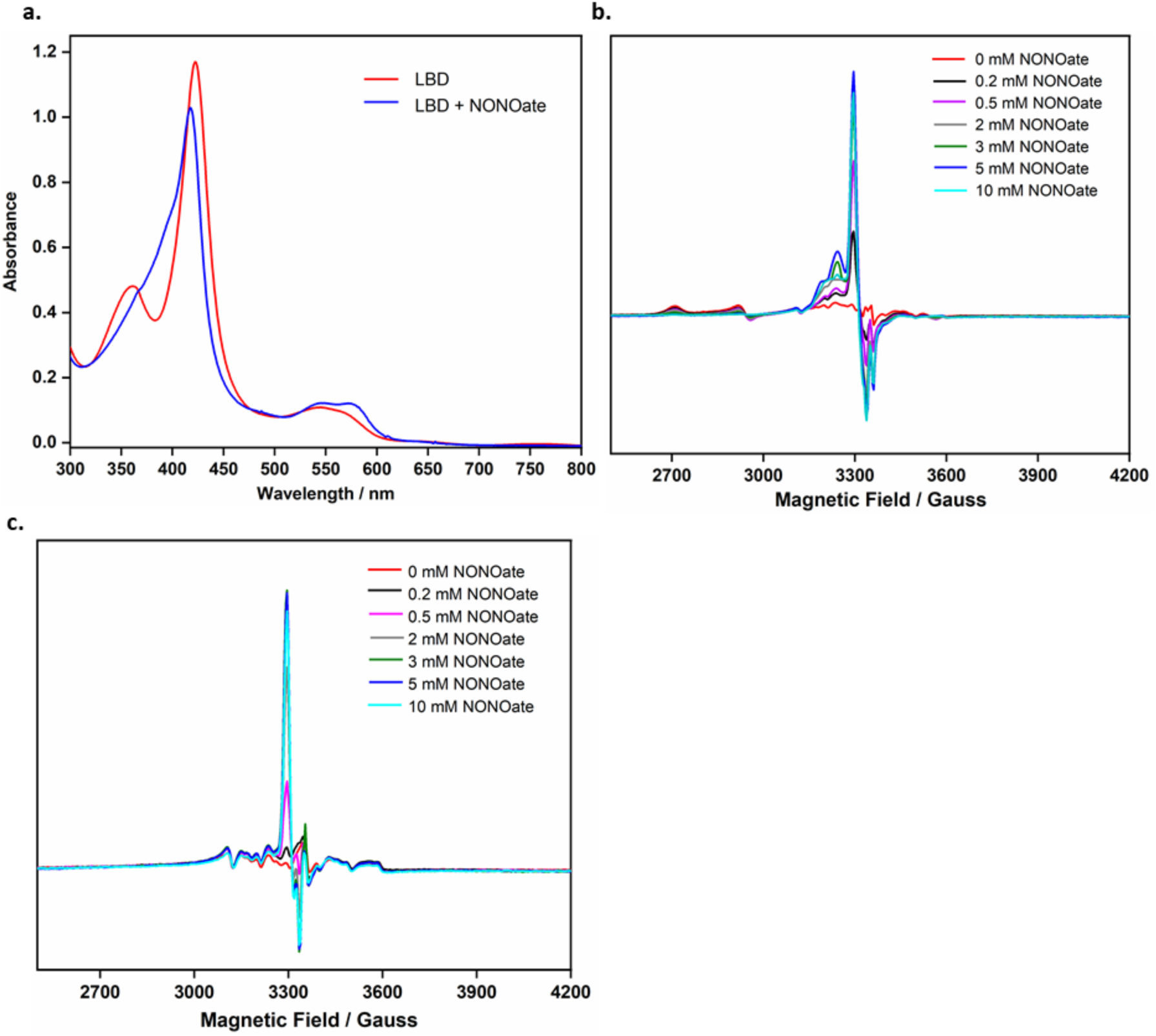
Effect of NO on EPR signal intensity of LBD heme. **a**, Absorption spectrum of the ferric LBD (93.2 µM), without redox mediator dyes, before and after Proline NONOate (2.5 mM) treatment; EPR spectra of *E. coli* cells **b**, over-expressing Rev-Erbβ, or **c**, containing empty vector pMCSG9, treated with increasing concentrations of Proline NONOate for 6 mins; EPR Conditions: temperature 11 K, microwave power 20 µW; microwave frequency 9.3831, 9.3815, 9.3828, 9.3817, 9.3823, 9.3831, 9.3823 GHz for **b** and 9.3857, 9.3842, 9.3831, 9.3829, 9.3813, 9.3832, 9.3840 GHz for **c**, modulation frequency 100 kHz, modulation amplitude 3 G, 4 scans, 327.68 ms time constant.

**Extended Data Fig. 6.**
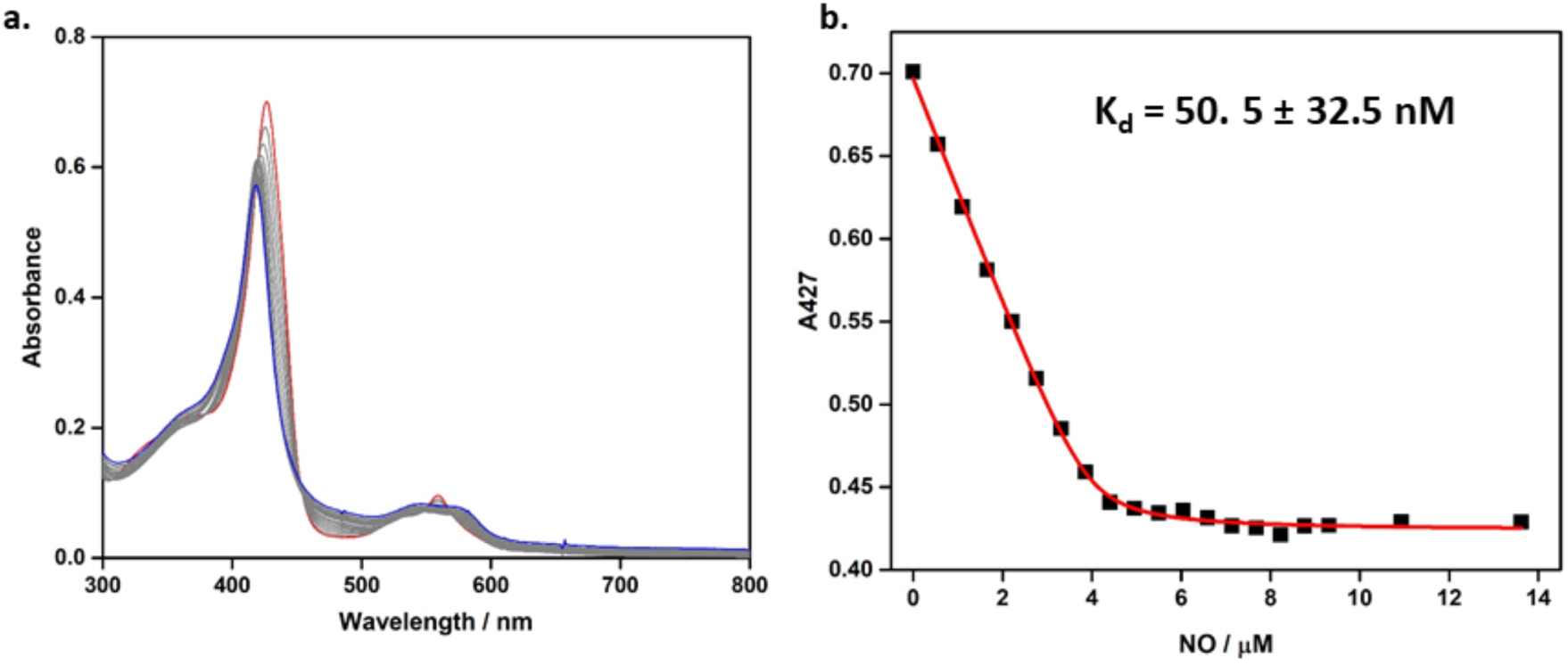
Binding Studies of Fe^2+^ Rev-Erbβ LBD with NO. **a**, Representative data set showing UV-Vis absorption change observed during titration of Fe^2+^ Rev-Erbβ with NO; **b**, Scatter plot of absorption at 427 nm versus NO concentration. Red line shows binding curve obtained by fitting the data to equation S3. K_d_ values reported here are an average of 5 experiments ± s.d.

**Extended Data Fig. 7.**
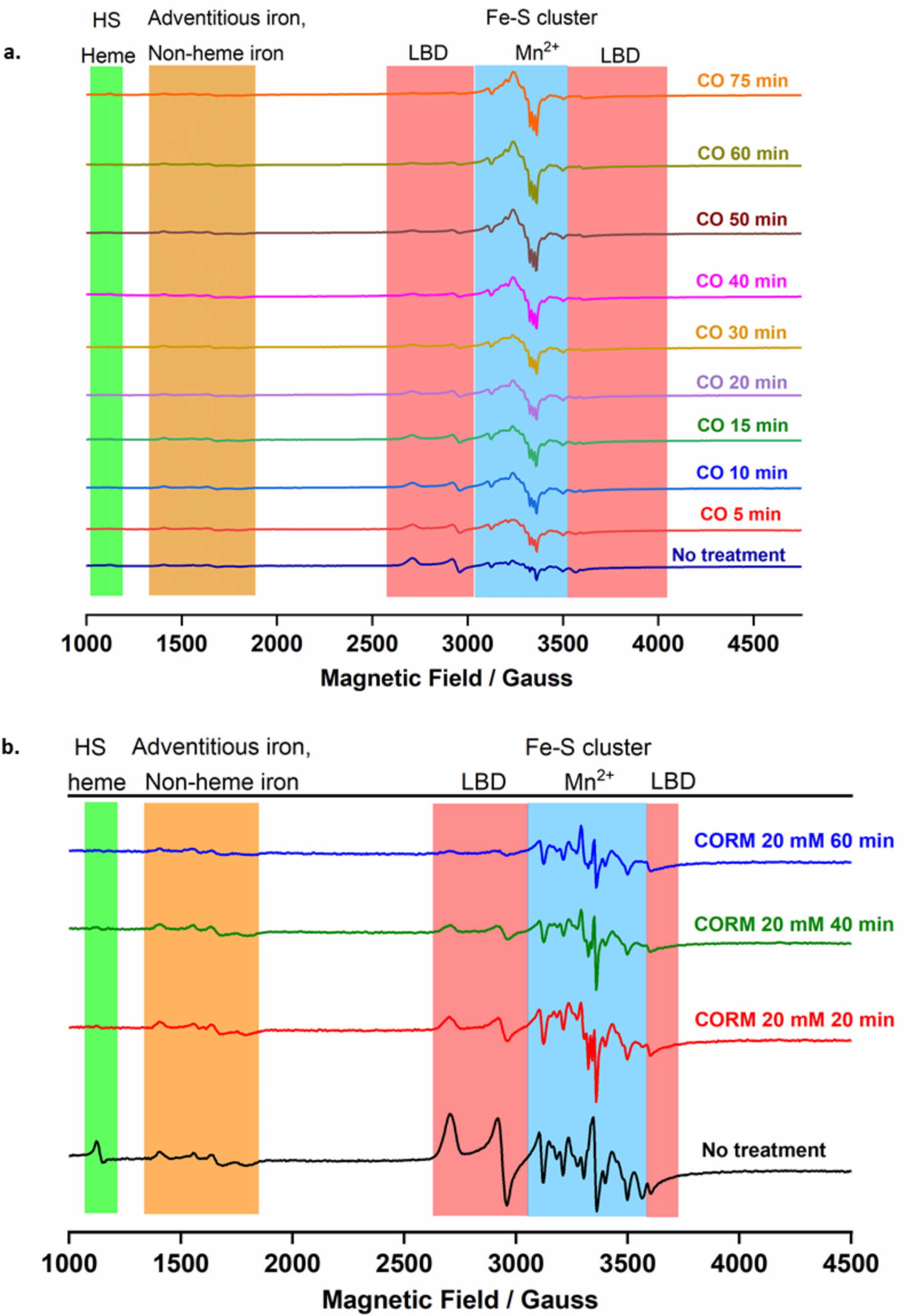
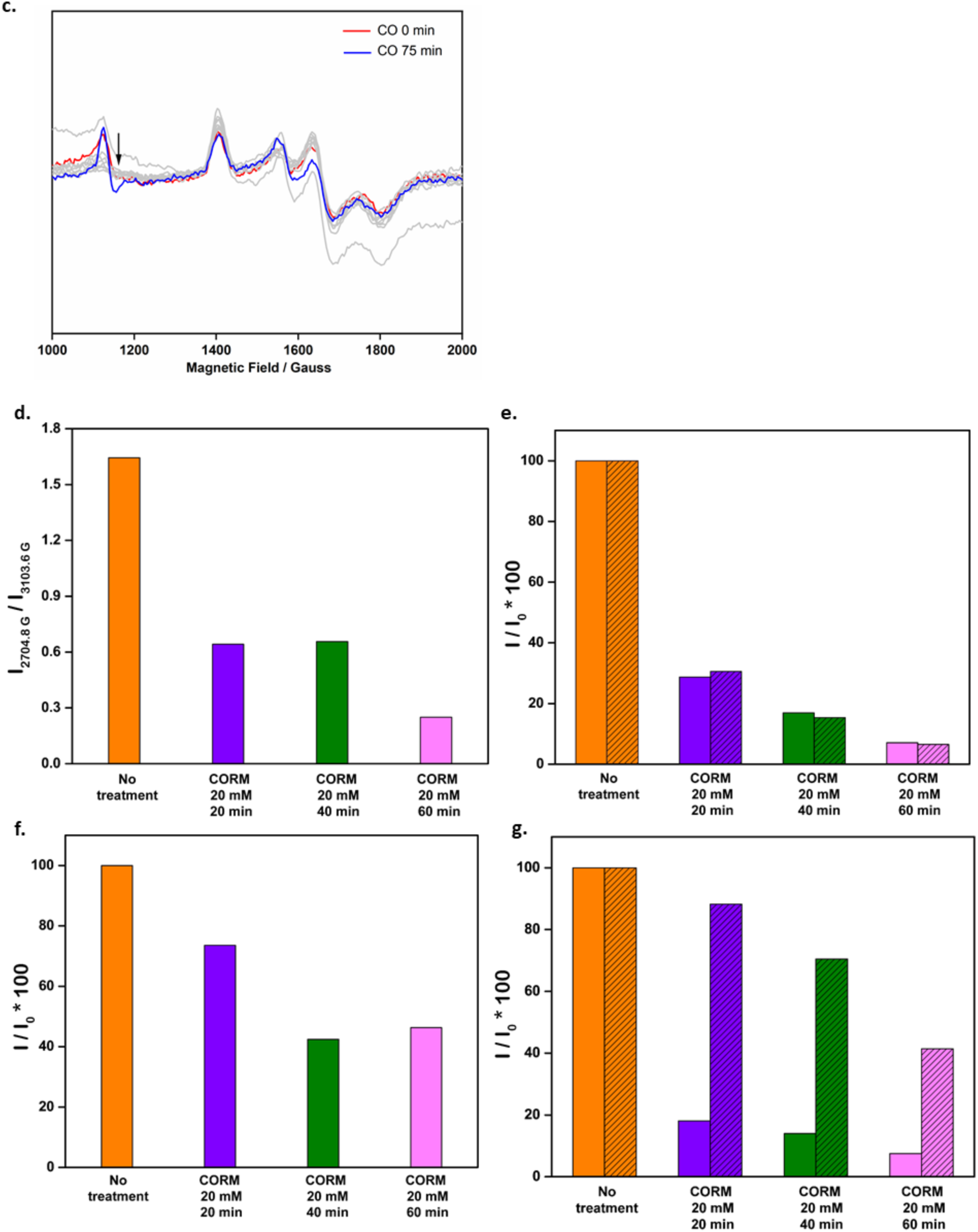
Effect of CO on the redox state of Rev-Erbβ *in cellulo*. Stacked EPR spectra of *E. coli* cells over-expressing Rev-Erbβ LBD, **a**, before and after CO purge for variable times, and **b**, with and without CORM-A1 treatment for variable times. The EPR spectra are divided into regions depicting spectral signature from LBD heme, Mn^2+^ and Fe-S Cluster, adventitious iron or non-heme iron, HS heme. **c**, Region depicting the EPR signal from adventitious iron, HS heme in **a. d**, Bar plots represent the ratio of the EPR signal intensity at 2704.8 G (characteristic of LBD heme) versus 3103.6 G (characteristic of Mn^2+^) from **b**; Bar plots represent percentage of observed EPR signal intensity with respect to the EPR signal intensity from *E. coli* cells without CORM treatment for the following field strengths: **e**, 2704.8 G (non-patterned bars) and 2919.84 G (patterned bars), characteristic of LBD heme; **f**, 3103.6 G, characteristic of cellular Mn^2+^; **g**, 1121.21 G (non-patterned bars) and 1402.7 G (patterned bars) characteristic of HS heme and non-heme iron respectively.

**Extended Data Fig. 8.**
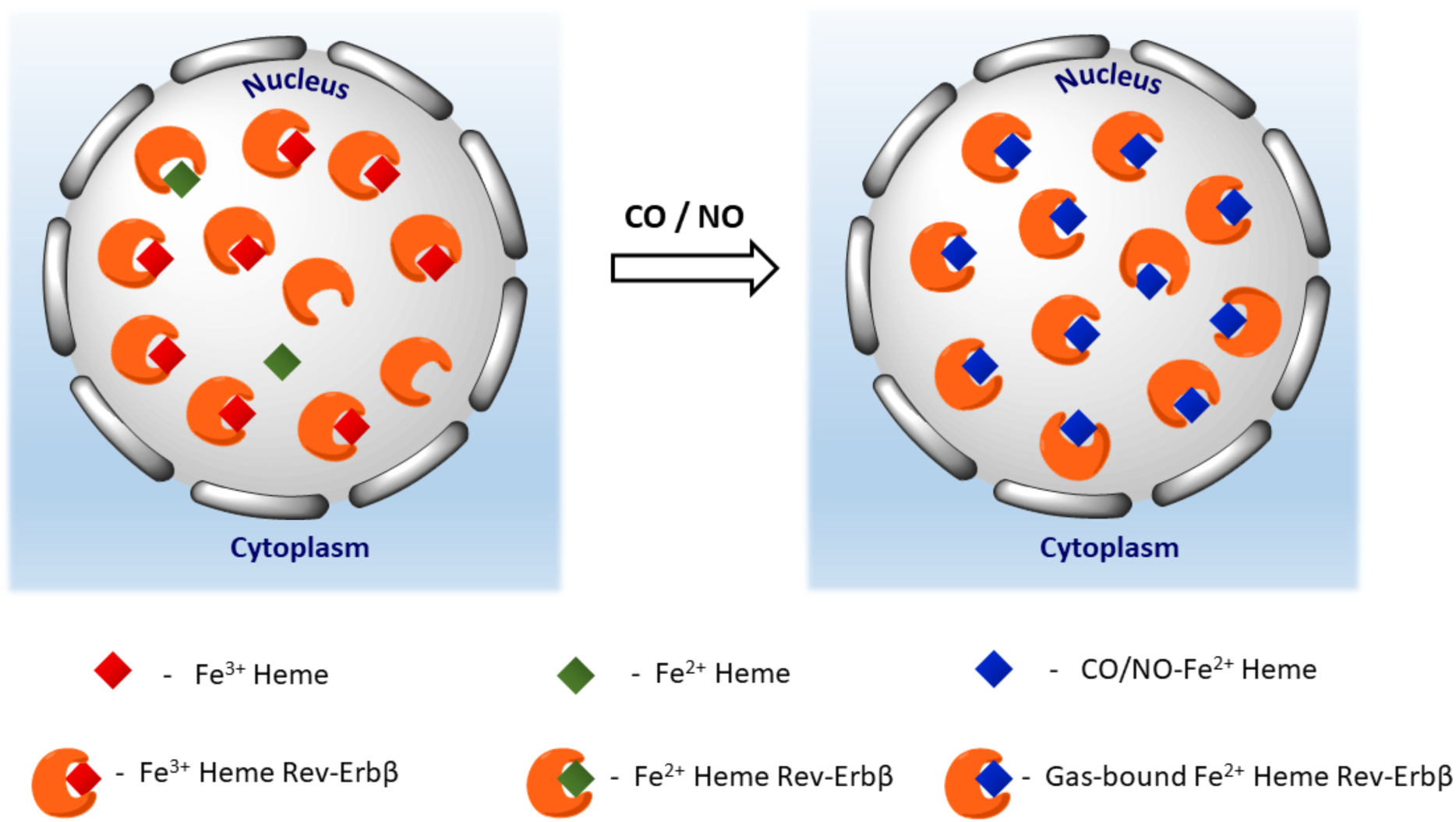
Rev-Erbβ exists in the Fe^3+^ heme-bound state in cells and can function as a gas sensor. Our experiments suggest that Rev-Erbβ predominantly exists Fe^3+^-heme bound state in cells. Under limiting concentrations of intracellular labile heme pools, Fe^3+^ not Fe^2+^ Rev-Erbβ remains in the heme-replete state. However, signaling gases like CO and NO exert a thermodynamic pull that allows the protein to be in the reduced gas-bound form. This intriguing redox chemistry allows the protein to shuffle between Fe^3+^ and Fe^2+^ heme states thereby not limiting Rev-Erb’s function as a gas-sensor even when the protein primarily exists in the oxidized form.

## Footnotes

1 However, the dithionite-reduced protein always shifted to 427 nm. EPR experiments ensured the integrity of the ferric protein (see Supplementary Fig. 1). Thus, we tentatively attribute the altered Soret absorption to an interaction between Rev-Erb and the redox mediators. Changes resulting from the datasets with no shift in the λmax of the Soret band for the ferrous protein are depicted in extended data Fig. 1b.

2 It has not yet been possible to obtain high enough concentrations of Rev-Erb to perform these experiments in human cells. However, the redox poise of the *E. coli* cytoplasm is similar to that of the nucleus of human cells in culture.^20^

3 Bindscheder’s Green was sparingly soluble in 0.5x TNG buffer and had to be handled anaerobically. Unlike others, the stock concentration of Bindscheder’s Green was less than 1 mM.

## References

1 Bugge, A. et al. Rev-erbalpha and Rev-erbbeta coordinately protect the circadian clock and normal metabolic function. Genes Dev 26, 657–667, (2012).

2 Cho, H. et al. Regulation of circadian behaviour and metabolism by REV-ERB-alpha and REV-ERB-beta. Nature 485, 123–127, (2012).

3 Crumbley, C. & Burris, T. P. Direct Regulation of CLOCK Expression by REV-ERB. PLOS ONE 6, e17290, (2011).

4 Ikeda, R. et al. REV-ERBα and REV-ERBβ function as key factors regulating Mammalian Circadian Output. Sci. Rep. 9, 10171, (2019).

5 Kojetin, D. J. & Burris, T. P. REV-ERB and ROR nuclear receptors as drug targets. Nat Rev Drug Discov 13, 197–216, (2014).

6 Kumar Jha, P., Challet, E. & Kalsbeek, A. Circadian rhythms in glucose and lipid metabolism in nocturnal and diurnal mammals. Mol. Cell. Endocrinol. 418, 74–88, (2015).

7 Lam, M. T. et al. Rev-Erbs repress macrophage gene expression by inhibiting enhancer-directed transcription. Nature 498, 511–515, (2013).

8 Carter, E. L., Gupta, N. & Ragsdale, S. W. High Affinity Heme Binding to a Heme Regulatory Motif on the Nuclear Receptor Rev-erbbeta Leads to Its Degradation and Indirectly Regulates Its Interaction with Nuclear Receptor Corepressor. J Biol Chem 291, 2196–2222, (2016).

9 Carter, E. L., Ramirez, Y. & Ragsdale, S. W. The heme-regulatory motif of nuclear receptor Rev-erbbeta is a key mediator of heme and redox signaling in circadian rhythm maintenance and metabolism. J Biol Chem 292, 11280–11299, (2017).

10 Raghuram, S. et al. Identification of heme as the ligand for the orphan nuclear receptors REV-ERBalpha and REV-ERBbeta. Nat Struct Mol Biol 14, 1207–1213, (2007).

11 Yin, L. et al. Rev-erbalpha, a heme sensor that coordinates metabolic and circadian pathways. Science 318, 1786–1789, (2007).

12 Gibbs, J. E. et al. The nuclear receptor REV-ERBα mediates circadian regulation of innate immunity through selective regulation of inflammatory cytokines. Proc. Natl. Acad. Sci. USA 109, 582–587, (2012).

13 Jager, J. et al. Behavioral Changes and Dopaminergic Dysregulation in Mice Lacking the Nuclear Receptor Rev-erbα. Mol. Endocrinol. 28, 490–498, (2014).

14 Schnell, A. et al. The Nuclear Receptor REV-ERBα Regulates Fabp7 and Modulates Adult Hippocampal Neurogenesis. PLOS ONE 9, e99883, (2014).

15 Valnegri, P. et al. A circadian clock in hippocampus is regulated by interaction between oligophrenin-1 and Rev-erbα. Nat. Neurosci. 14, 1293, (2011).

16 Carter, E. L. & Ragsdale, S. W. Modulation of nuclear receptor function by cellular redox poise. J Inorg Biochem 133, 92–103, (2014).

17 Shimizu, T. et al. Gaseous O2, NO, and CO in Signal Transduction: Structure and Function Relationships of Heme-Based Gas Sensors and Heme-Redox Sensors. Chem. Rev. 115, 6491–6533, (2015).

18 Marvin, K. A. et al. Nuclear Receptors Homo sapiens Rev-erbβ and Drosophila melanogaster E75 Are Thiolate-Ligated Heme Proteins Which Undergo Redox-Mediated Ligand Switching and Bind CO and NO. Biochemistry 48, 7056–7071, (2009).

19 Pardee, K. I. et al. The Structural Basis of Gas-Responsive Transcription by the Human Nuclear Hormone Receptor REV-ERBβ. PLoS Biol. 7, e1000043, (2009).

20 Gupta, N. & Ragsdale, S. W. Thiol-disulfide redox dependence of heme binding and heme ligand switching in nuclear hormone receptor rev-erb{beta}. J Biol Chem 286, 4392–4403, (2011).

21 Hanna, D. A. et al. Heme dynamics and trafficking factors revealed by genetically encoded fluorescent heme sensors. Proc Natl Acad Sci U S A 113, 7539–7544, (2016).

22 Caceres, L. et al. Nitric oxide coordinates metabolism, growth, and development via the nuclear receptor E75. Genes Dev 25, 1476–1485, (2011).

23 Reinking, J. et al. The Drosophila nuclear receptor e75 contains heme and is gas responsive. Cell 122, 195–207, (2005).

24 Go, Y. M. & Jones, D. P. Redox control systems in the nucleus: mechanisms and functions. Antioxid Redox Signal 13, 489–509, (2010).

25 Reedy, C. J., Elvekrog, M. M. & Gibney, B. R. Development of a heme protein structure– electrochemical function database. Nucleic Acids Res. 36, D307–D313, (2007).

26 Bickar, D., Bonaventura, C. & Bonaventura, J. Carbon monoxide-driven reduction of ferric heme and heme proteins. J Biol Chem 259, 10777–10783, (1984).

27 Can, M., Armstrong, F. A. & Ragsdale, S. W. Structure, function, and mechanism of the nickel metalloenzymes, CO dehydrogenase, and acetyl-CoA synthase. Chem Rev 114, 4149–4174, (2014).

28 Hirota, S., Azuma, K., Fukuba, M., Kuroiwa, S. & Funasaki, N. Heme reduction by intramolecular electron transfer in cysteine mutant myoglobin under carbon monoxide atmosphere. Biochemistry 44, 10322–10327, (2005).

29 Ford, P. C., Fernandez, B. O. & Lim, M. D. Mechanisms of Reductive Nitrosylation in Iron and Copper Models Relevant to Biological Systems. Chem. Rev. 105, 2439–2456, (2005).

30 Silkstone, G., Kapetanaki, S. M., Husu, I., Vos, M. H. & Wilson, M. T. Nitric oxide binds to the proximal heme coordination site of the ferrocytochrome c/cardiolipin complex: formation mechanism and dynamics. J Biol Chem 285, 19785–19792, (2010).

31 Wareham, L. K., Southam, H. M. & Poole, R. K. Do nitric oxide, carbon monoxide and hydrogen sulfide really qualify as ‘gasotransmitters’ in bacteria? Biochemical Society Transactions 46, 1107–1118, (2018).

32 Tinberg, C. E. et al. Characterization of iron dinitrosyl species formed in the reaction of nitric oxide with a biological Rieske center. J. Am. Chem. Soc. 132, 18168–18176, (2010).

33 Carballal, S. et al. Kinetics of reversible reductive carbonylation of heme in human cystathionine beta-synthase. Biochemistry 52, 4553–4562, (2013).

34 Hammes-Schiffer, S. & Stuchebrukhov, A. A. Theory of coupled electron and proton transfer reactions. Chem. Rev. 110, 6939–6960, (2010).

35 Leferink, N. G. et al. Gating mechanisms for biological electron transfer: integrating structure with biophysics reveals the nature of redox control in cytochrome P450 reductase and copper-dependent nitrite reductase. FEBS Lett 586, 578–584, (2012).

36 Johnston, W. A. et al. Cytochrome P450 is present in both ferrous and ferric forms in the resting state within intact Escherichia coli and hepatocytes. J Biol Chem 286, 40750–40759, (2011).

37 Kapetanaki, S. M. et al. A mechanism for CO regulation of ion channels. Nat. Commun. 9, 907, (2018).

38 Nakajima, H. et al. Redox properties and coordination structure of the heme in the co-sensing transcriptional activator CooA. J Biol Chem 276, 7055–7061, (2001).

39 Carballal, S. et al. Dioxygen Reactivity and Heme Redox Potential of Truncated Human Cystathionine β-Synthase. Biochemistry 47, 3194–3201, (2008).

40 Ding, X. D. et al. Nitric Oxide Binding to the Ferri- and Ferroheme States of Nitrophorin 1, a Reversible NO-Binding Heme Protein from the Saliva of the Blood-Sucking Insect, Rhodnius prolixus. J. Am. Chem. Soc. 121, 128–138, (1999).

## Methods References

41 Barr, I. & Guo, F. Pyridine Hemochromagen Assay for Determining the Concentration of Heme in Purified Protein Solutions. Bio. Protoc. 5, e1594, (2015).

42 Dawson, R. M. C., Elliott, D. C., Elliott, W. H. & Jones, K. M. Data for biochemical research. (2nd edn). (Clarendon Press, 1969).

43 Homer, R. F. & Tomlinson, T. E. 504. The stereochemistry of the bridged quaternary salts of 2,2&-bipyridyl. J. Chem. Soc. (Resumed), 2498–2503, (1960).

44 Efimov, I. et al. A simple method for the determination of reduction potentials in heme proteins. FEBS Lett. 588, 701–704, (2014).

45 Clark, W. M. Oxidation-reduction potentials of organic systems. (Robert E. Krieger Publishing Company, 1972).

46 Salmon, R. T. & Hawkridge, F. M. The electrochemical properties of three dipyridinium salts as mediators. J. Electroanal. Chem. Interf. Electrochem. 112, 253–264, (1980).

47 Wardman, P. Reduction Potentials of One-Electron Couples Involving Free Radicals in Aqueous Solution. J. Phys. Chem. Ref. Data 18, 1637–1755, (1989).

48 Chan, M. S. & Bolton, J. R. Structures, reduction potentials and absorption maxima of synthetic dyes of interest in photochemical solar-energy storage studies. Sol. Energy 24, 561–574, (1980).

