## Supplemental Fig. 1 for "Ferric Heme as a CO/NO Sensor in the Nuclear Receptor Reverbβ by Coupling Gas binding to Electron Transfer"

**Thermodynamic box calculations**

Derivation of the thermodynamic parameters for different equilibria depicted in the thermodynamic box in Fig. 2d, are given below. [red] and [oxd] indicate the concentration of reduced and oxidized protein.

*I. $Fe^{3+/2+}$ redox equilibrium*

$$Fe^{3+}\ LBD \rightleftharpoons Fe^{2+}\ LBD$$

Our redox titrations yield two populations of the proteins with E° values of - 0.225 V and – 0.405 V and relative amplitudes of 0.4136 and 0.5864, respectively (extended data table 1). The relative content of the reduced and oxidized forms for each population at a potential of -280 mV (which is the nuclear potential)[9] was calculated based on the Nernst equation (S4).

$$E = E^0 - 0.0591 log\frac{[red]}{[oxd]} \quad \ldots\ldots\ldots\ldots S4$$

Taking into account the amplitudes of each of these populations and the individual ratios of oxidized versus reduced protein, overall 62.5% of the protein exists in the oxidized and 37.5% of the protein in the reduced form.

$$\frac{[oxd]}{[red]} = \frac{0.625}{0.375}$$

*II. $Fe^{3+/2+}$ redox equilibrium in the presence of CO/NO*

$$Fe^{3+}\text{-g LBD} \rightleftharpoons Fe^{2+}\text{-g LBD} \qquad \text{where g is CO/NO}$$

Our redox titrations indicate the E° for $Fe^{3+}$-g/$Fe^{2+}$-g equilibrium > 989 mV for CO and > 819 mV for NO. The ratio of the reduced to oxidized protein in the presence of the gases at a potential of -280 mV calculated using equation S4 is

$\frac{[oxd]}{[red]} > 10^{21.5}$ for CO and

$\frac{[oxd]}{[red]} > 10^{18.6}$ for NO.

Therefore, for our redox reaction with CO (Fig. 2b) when the total concentration of Rev-Erbβ LBD protein was 95.5 µM,

[oxd] < $9.55 \times 10^{-20.5}$ µM and [red] ≅ 95.5 – $9.55 \times 10^{-20.5}$ ≅ 95.5 µM.

Similarly, for redox titrations in presence of NO (Fig. 2c) when the total concentration of LBD protein was 99.1 µM,

[oxd] < $9.91 \times 10^{-18.6}$ µM and [red] ≅ 99.1 – $9.91 \times 10^{-18.6}$ ≅ 99.1 µM.

*III. Dissociation Constant of $Fe^{3+}$-g LBD*

$$Fe^{3+}LBD + g \rightleftharpoons Fe^{3+}LBD\text{-}g \qquad \text{where g is CO/NO}$$

Dissociation constant, $K_{d2}$, for the above reaction is given by

$$K_{d2} = \frac{[Fe^{3+}LBD]\times[g]}{[Fe^{3+}LBD-g]} \text{........S5}$$

For the redox titrations in presence of CO,
$[Fe^{3+}LBD] \cong 95.5 - 9.55 \times 10^{-20.5} \cong 95.5$ μM; [CO] = 439 μM (determined as mentioned previously); $[Fe^{3+}LBD\text{-}CO] < 9.55 \times 10^{-20.5}$ μM.

Therefore, $K_{d2}$ (in presence of CO) > $4.39 \times 10^{17.5}$ M

For the redox titrations in presence of NO,
$[Fe^{3+}LBD] \cong 99.1 - 9.91 \times 10^{-18.6} \cong 99.1$ μM; [NO] = 1.94 mM (solubility of NO in water)[10]; $[Fe^{3+}LBD\text{-}NO] < 9.91 \times 10^{-18.6}$ μM.

Therefore, $K_{d2}$ (in presence of NO) > $1.94 \times 10^{16.6}$ M

**Control experiments to explain the origin of 422 nm Soret peak of the reduced protein**

We performed a few control experiments to account for the serendipitous occurrence of the 422 nm Soret peak for the reduced protein.

1. Integrity of the $Fe^{3+}$ protein was checked by EPR spectroscopy. The EPR indicated the absence of any form of LS or HS ferric heme other than the His/Cys-LS form (Supplemental fig 1).
2. The addition of dye cocktail did not elicit any significant spectral change in the protein eliminating non-specific interaction of the ferric protein with dyes (Supplemental fig 2a).
3. The chemical reduction of the protein (with dyes) using the dithionite yielded the reduced protein with Soret peak at 427 nm eliminating non-specific interaction of the reduced protein with dyes (Supplemental fig 2b).
4. The integrity of the CV-27 voltammogram was verified by using a multimeter in parallel to the electrode set-up in the cuvette to re-confirm the values of applied potential.

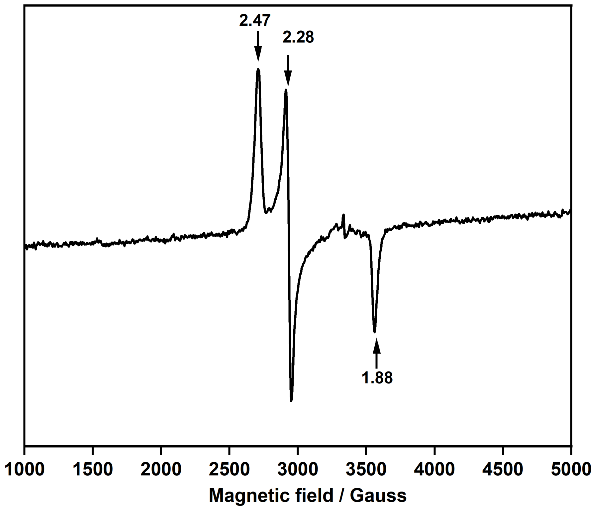


**Supplementary Fig. 1 Spectroscopic evidence of Rev-Erbβ LBD with His/Cys coordinated $Fe^{3+}$-heme.** EPR spectrum of Rev-Erbβ LBD depicting the rhombic spectra with *g*-values indicating only His/Cys coordinated LS-$Fe^{3+}$ heme. EPR Conditions: temperature 10 K, microwave power 206 μW; microwave frequency 9.2680 GHz, modulation frequency 100 kHz, modulation amplitude 7 G, 2 scans, 327.68 ms time constant.

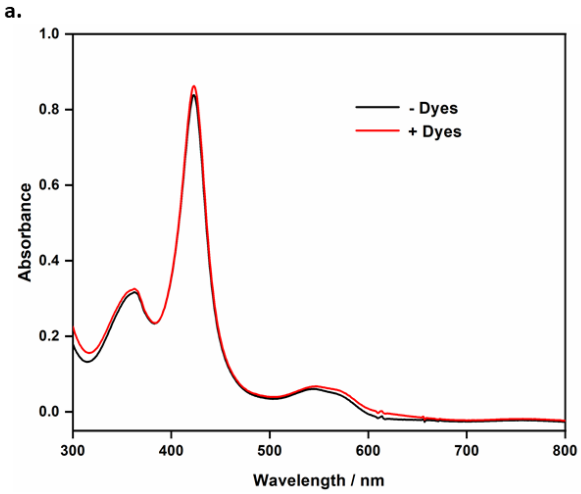


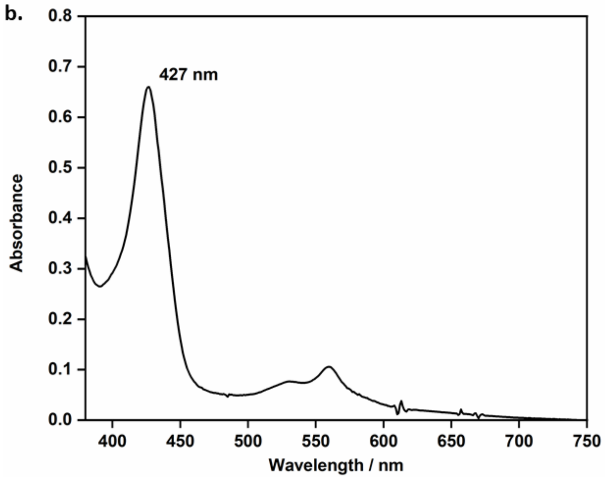


**Supplementary Fig. 2 Interaction of dyes with the $Fe^{3+}$ and $Fe^{2+}$-LBD heme. a,** Absorption spectrum of the $Fe^{3+}$-LBD heme with and without dyes; **b,** Absorption spectrum of the $Fe^{3+}$-LBD protein containing mediator dyes post treatment with dithionite.
